# Accurate strand-specific long-read transcript isoform discovery and quantification at bulk, single-cell, and single-nucleus resolution

**DOI:** 10.64898/2026.02.12.705617

**Authors:** Houlin Yu, Christophe H. Georgescu, Akanksha Khorgade, Ghamdan Al-Eryani, Daniel A. Bartlett, Allison Brookhart, Can Kockan, James T. Webber, Asa Shin, Emily White, Xylena Reed, Fangle Hu, Sarah Bromberek, Sandra Ndayambaje, Sandeep Aryal, Dennis W. Dickson, Mercedes Prudencio, Clotilde Lagier-Tourenne, Michael E. Ward, Paul C. Blainey, Victoria Popic, Brian J. Haas, Aziz M. Al’Khafaji

## Abstract

Recent advances in long-read transcriptome sequencing enable high-throughput profiling of full-length RNA isoforms in bulk, single-cell, and single-nucleus samples. However, long-read datasets typically contain a mixture of complete and partial transcripts, leading to pervasive ambiguity in read-to-isoform assignment and complicating accurate isoform identification and quantification, particularly in the absence of reliable reference annotations. These challenges are further amplified in single-cell and single-nucleus samples, where coverage is sparse and transcriptional heterogeneity is high.

Here, we present the Long Read Alignment Assembler (LRAA), a unified and versatile computational framework for isoform identification and quantification from long-read RNA sequencing data across bulk, single-cell, and single-nucleus transcriptomic samples. LRAA combines splice-graph based structural modeling with expectation maximization based optimization to probabilistically resolve ambiguous read assignments and improve isoform abundance estimation. The framework supports quantification-only, reference-guided, and fully reference-free (de novo) modes of analysis within a single methodological paradigm.

We benchmarked LRAA using both simulated and genuine long-read datasets spanning sequencing standards and whole transcriptomes. Central to this evaluation is a novel benchmarking strategy based on Multiplexed Overexpression of Regulatory Factors (MORFs), which provides biologically expressed, barcoded isoforms with unambiguous read-level ground truth. Across all benchmarks, including MORFs, synthetic spike-ins, and whole-transcriptome datasets, LRAA consistently outperformed state-of-the-art methods in isoform identification accuracy, sensitivity, and expression quantification.

Finally, we demonstrate the biological utility of LRAA by resolving cell-type-specific isoform usage across peripheral blood immune cell populations and by detecting a pathogenic cryptic isoform of *STMN2* with associated transcriptional changes in single-nucleus RNA-seq data from frontal cortex tissue of an individual with frontotemporal dementia (FTD). Together, these results establish LRAA as a robust and general solution for resolving transcript diversity in complex biological systems, from development to disease.

## Introduction

Transcriptome sequencing is a foundational biological assay that enables detailed phenotypic characterization by measuring gene expression, RNA splicing, and transcript diversity. This technology provides critical insights into cellular state and function across various tissues and biological contexts, including health, development, and disease (Stark et al., 2019; Z. Wang et al., 2009). Accurate resolution of transcript isoforms is essential for understanding gene regulation and function, as the vast majority of genes undergo alternative splicing, producing multiple RNA isoforms with distinct biological functions (Keren et al., 2010; Nilsen & Graveley, 2010). Additionally, transcriptome profiling also underpins genome annotation, particularly in complex tissues and non-model organisms where reference annotations remain incomplete (Grabherr et al., 2011; Haas et al., 2002; M. Pertea et al., 2018).

Historically, transcriptomics has relied on short-read sequencing technologies, which offer high throughput and accurate gene expression quantification but provide only fragmented views of RNA molecules (Mortazavi et al., 2008). Reconstruction of full-length transcripts from short reads requires computational assembly or inference, which is inherently ambiguous for genes with complex splicing patterns, alternative transcription start sites, or alternative polyadenylation (Garber et al., 2011; Martin & Wang, 2011). As a result, accurate resolution of transcript structure has remained a persistent bottleneck, limiting both isoform discovery and reliable quantification (Steijger et al., 2013; Teng et al., 2016).

Long-read sequencing technologies, including platforms from Pacific Biosciences and Oxford Nanopore Technologies, have fundamentally changed this landscape by enabling sequencing of full-length RNA molecules (Sharon et al., 2013; Tilgner et al., 2014). These approaches allow transcript structures to be observed directly, reducing reliance on computational reconstruction and revealing extensive isoform diversity that is systematically missed by short-read methods. Recent improvements in throughput, accuracy, and cost effectiveness, including high-throughput long-read protocols applicable to bulk, single-cell, and single-nucleus RNA sequencing, have made isoform-resolved transcriptomics broadly accessible across biological and clinical applications (Al’Khafaji et al., 2024; Amarasinghe et al., 2020; Volden et al., 2018).

Despite these advances, computational analysis remains a major limiting factor. Long-read RNA sequencing datasets frequently contain partial or truncated transcripts, heterogeneous error profiles, and extensive ambiguity in read-to-isoform compatibility, particularly for genes with shared exons or incomplete coverage (Amarasinghe et al., 2020; Tardaguila et al., 2018). These challenges are compounded in single-cell and single-nucleus settings, where coverage is sparse and transcriptional heterogeneity is high (I. Gupta et al., 2018; Hardwick et al., 2022). As a result, accurate isoform identification and quantification from long-read data remains an unsolved problem, particularly in contexts where reference annotations are incomplete or biased.

To address these challenges, a diverse ecosystem of computational tools for long-read transcriptome analysis has emerged, reflecting different design priorities and trade-offs (Amarasinghe et al., 2020). Several methods emphasize accurate reconstruction of transcript structure from genome-aligned long reads. Tools such as FLAIR (Tang et al., 2020), TALON (Wyman et al., 2019), Mandalorion (Volden et al., 2023), ESPRESSO (Gao et al., 2023), and IsoQuant (Prjibelski et al., 2023) have demonstrated strong performance for isoform identification, often prioritizing precision through conservative modeling, explicit splice junction validation, or robust handling of sequencing errors. These approaches have been instrumental in establishing reliable long-read isoform catalogs, particularly in reference-guided or high-coverage settings (Glinos et al., 2022; Reese et al., 2023; Workman et al., 2018, 2019). Other methods have focused on transcript abundance estimation in the presence of ambiguous read-to-isoform compatibility. Probabilistic frameworks based on expectation maximization, including Oarfish (Zare Jousheghani et al., 2025) and Isosceles (Kabza et al., 2024), explicitly model uncertainty in read assignment and can achieve accurate quantification when transcript models are known or partially reconstructed. Similarly, Bambu (Chen et al., 2023) integrates reference-guided transcript discovery with statistical modeling to improve quantification under incomplete annotations. The capabilities and scope of representative long-read isoform analysis methods, including support for bulk, single-cell, and single-nucleus data, are summarized in **Table 1**. While these methods provide important advances in abundance estimation, they typically assume an externally defined set of isoforms and do not fully integrate de novo isoform discovery and quantification within a single inference framework.

**Table 1.** Capabilities of Related Methods for Long Read-based Isoform Identification and Quantification.

| Tools | Reference-guided Identification | *De novo* Identification | Expression Quantification | Single-cell Transcriptome Applications | Reference |
| --- | --- | --- | --- | --- | --- |
| LRAA | ✓ | ✓ | ✓ | ✓ | This study |
| IsoSeq | ✓ | ✓ | ✓ | ✓ | (*Iso-Seq Home*, n.d.) |
| IsoQuant | ✓ | ✓ | ✓ | ** | (Prjibelski et al., 2023) |
| Bambu | ✓ | ✓ | ✓ | ** | (Chen et al., 2023) |
| Mandalorion | ✓ | ✓ | ✓ |  | (Volden et al., 2023) |
| StringTie | ✓ | ✓ | ✓ |  | (Kovaka et al., 2019) |
| Isosceles | ✓ |  | ✓ | ✓ | (Kabza et al., 2024) |
| ESPRESSO | ✓ |  | ✓ | * | (Gao et al., 2023) |
| FLAMES | ✓ |  | ✓ | ** | (Holmqvist et al., 2021) |
| FLAIR | ✓ |  | ✓ |  | (Tang et al., 2020) |
| TALON | ✓ |  | ✓ |  | (Wyman et al., 2019) |
| Oarfish |  |  | ✓ |  | (Zare Jousheghani et al., 2025) |

\* read to isoform assignments can allow for cell barcode identification \*\*recently developed Bambu-Clump (Sim et al., 2025), Flames-v2 (C. Wang et al., 2025), and Spl-IsoQuant-2 (Michielsen et al., 2025) support single-cell and/or spatial long-read transcriptomics were not evaluated here.

As a result, existing tools tend to excel either at isoform structure reconstruction or at probabilistic abundance estimation but rarely address both problems simultaneously within a unified framework, particularly in annotation-free settings and in datasets dominated by partial or truncated transcripts. This separation becomes especially limiting for single-cell and single-nucleus long-read transcriptomics, where sparse coverage, heterogeneous transcript boundaries, and extensive read ambiguity further exacerbate the coupling between isoform discovery and quantification.

Rigorous benchmarking of long-read isoform identification and quantification methods is another major challenge. A recent community-led effort, the Long-read RNA-Seq Genome Annotation Assessment Project (LRGASP), evaluated the accuracy of computational approaches for isoform identification and quantification across a broad range of methods, datasets, and long-read sequencing protocols (Pardo-Palacios, Wang, et al., 2024). This study revealed substantial variability in performance across tools, with no single method consistently achieving strong results across all evaluation settings, highlighting both progress in the field and continued room for methodological improvement. Importantly, LRGASP also underscored limitations of existing benchmarking substrates, particularly those relying exclusively on simulated data or on genuine transcriptomes lacking definitive read-level ground truth, motivating the development of additional benchmarking datasets and evaluation strategies.

An ideal benchmark would combine well-defined transcript structures, expression within a native cellular context, and unambiguous read-level ground truth. However, no commonly used benchmarking strategy satisfies all three criteria. Synthetic spike-in controls, such as SIRVs and Sequins, provide predefined transcript structures but lack endogenous transcriptional regulation and RNA processing, and discrepancies between expected and observed abundances can conflate experimental variability with computational performance. Conversely, benchmarks derived from biological samples preserve native cellular context but lack definitive isoform-level ground truth, often relying on proxy evaluations such as transcript hold-outs or cross-method consensus that do not reflect real-world discovery scenarios within annotated genes (Pardo-Palacios, Wang, et al., 2024; Steijger et al., 2013).

Critically, existing benchmarks rarely provide read-level truth assignments for partial or ambiguous reads, despite such reads being pervasive in long-read RNA sequencing datasets. This limitation hinders principled assessment of both isoform reconstruction and quantification accuracy, particularly in complex genes where multiple isoforms share substantial sequence overlap. Together, these considerations underscore the need for benchmarking frameworks that combine biological realism with absolute, read-resolved ground truth.

In this work, we address both methodological and benchmarking gaps. First, we introduce the Long Read Alignment Assembler (LRAA), a unified framework for isoform identification and quantification from long-read RNA sequencing data. LRAA combines splice-graph–based structural modeling with an expectation-maximization optimization framework, enabling probabilistic resolution of ambiguous read assignments while retaining sensitivity to novel isoforms. Second, to support rigorous evaluation under biologically realistic conditions, we introduce a benchmarking strategy based on Multiplexed Overexpression of Regulatory Factors (MORFs) (Joung et al., 2023), which yields biologically expressed, barcoded isoforms with unambiguous read-level ground truth.

We apply LRAA across simulated and genuine long-read datasets (**Supplementary Table 1**) spanning bulk, single-cell, and single-nucleus transcriptomes, and demonstrate its utility in resolving cell-type-specific isoform usage in peripheral blood immune cells as well as disease-associated cryptic splicing in neuronal cells of a human frontal cortex tissue sample from an individual with frontotemporal dementia (FTD).

## Results

### Long Read Alignment Assembler (LRAA) for isoform identification and quantification

LRAA was designed as a unified framework for reference-guided and reference-free isoform discovery and quantification from spliced genome alignments of long-read RNA sequencing data. The method explicitly addresses two pervasive challenges in long-read transcriptomics: incomplete transcript coverage that yields partial isoform fragments, and ambiguity in read compatibility across overlapping transcript isoform structures. To resolve these challenges, LRAA integrates splice-graph based structural modeling with probabilistic isoform quantification using expectation maximization (EM), allowing isoform discovery and abundance estimation to be tightly coupled within a single inference framework (**Fig. 1** and **Supplementary Fig. 1**).

**Figure 1:**
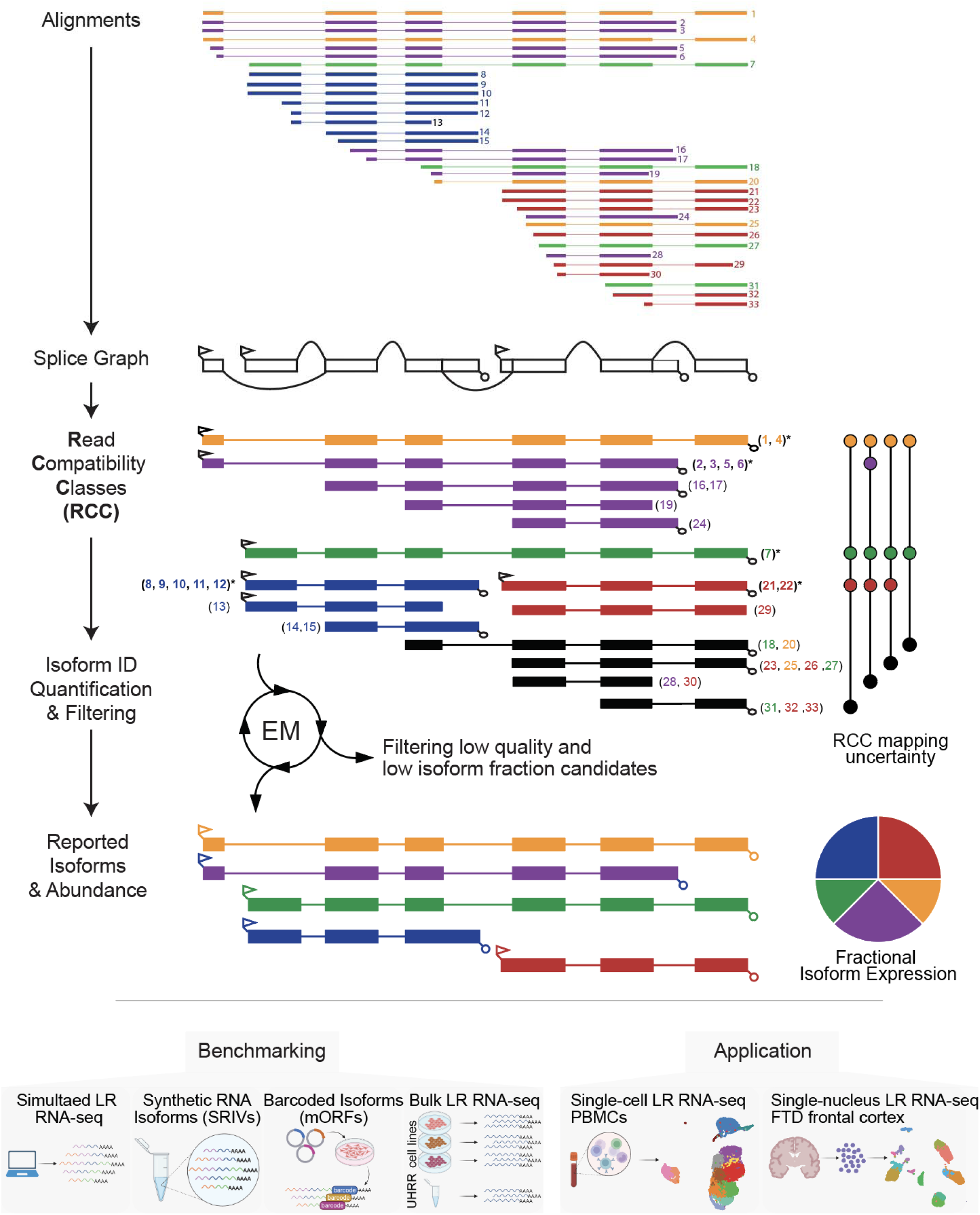
Conceptual overview of the LRAA algorithm for isoform identification and quantification in bulk and single-cell long-read transcriptomics. A simplified overview of the LRAA workflow is shown, beginning with splice-aligned long-read isoform data and encompassing splice-graph construction, assignment of read alignments to path nodes and grouping into read compatibility classes (RCC), assembly of candidate isoforms via compatible RCC collapse, and iterative quantification and filtering to define final isoform structures and abundance estimates. For illustrative purposes, read alignments are colored according to the isoforms from which they were derived and numbered to enable visual tracking through the splice graph and RCC relationships; in real RNA-seq data, the transcript of origin for each read is unknown a priori and must be fully inferred from alignment structures and compatibility relationships. Reads are numbered to allow tracking to the splice graph-based RCCs to which they define. Top-level RCCs not compatible and contained by other RCCs are labeled with *. RCCs found compatible and contained by other top-level RCCs are colored black and the RCC mapping uncertainty matrix indicates the multiple compatibilities. RCCs with top-level RCC mapping uncertainty have read support fractionally assigned according to the EM. For clarity, several implementation details are abstracted in this schematic; a complete description of the algorithmic workflow is provided in Supplementary Figure 1, which illustrates the construction of separate splice graphs for multi-exon and single-exon read alignments, their subsequent merging, and the downstream quantification and filtering steps. In the schematic, 5′ pointed triangles denote candidate transcription start sites (TSS), and 3′ trailing lollipops denote candidate transcription end sites (TES) or sites of polyadenylation. The LRAA identified isoforms were leveraged for multiple benchmarking and applications as shown (bottom).

LRAA operates on strand-specific spliced alignments of long reads to the reference genome. For each chromosome and transcriptional orientation, LRAA constructs a splice graph whose nodes represent partitioned exon segments, splice junctions, and transcript boundary features including transcription start sites (TSS) and transcription end sites (TES), typically corresponding to polyadenylation sites. Separate splice graphs are constructed for multi-exonic and single-exon alignments to prevent inappropriate coupling of spliced and unspliced read structures (**Supplementary Fig. 1**).

Each read alignment is decomposed into its corresponding path through the splice graph and encoded as an ordered sequence of labeled graph elements. Read alignments are grouped into read compatibility classes (RCCs), which represent sets of reads that are structurally indistinguishable with respect to exon–intron architecture and transcript boundaries according to paths traversed in the splice graph. This representation provides a compact description of read support while explicitly capturing ambiguity arising from partial transcript coverage and shared exons.

Candidate isoforms are assembled by hierarchically merging transcriptionally compatible RCCs based on shared splice patterns, transcript boundaries, and genomic overlap. Smaller RCCs that are fully compatible with larger mutually exclusive RCCs may be subsumed during this process, reflecting cases in which partial reads are consistent with multiple longer transcript models. Assembled isoforms are grouped into gene-level components based on shared splice graph structure and overlap thresholds, yielding a set of candidate isoforms and their associated RCC support.

Isoform abundance estimation is performed independently within each gene-level component using an EM algorithm that fractionally assigns reads from shared RCCs to compatible isoforms based on maximum likelihood. Reads that are uniquely compatible with a single isoform contribute entirely to its abundance estimate, whereas reads compatible with multiple isoforms are probabilistically allocated according to current isoform proportion estimates. To mitigate over-fragmentation of isoform models and suppress low-confidence transcripts, LRAA applies iterative filtering during quantification.

Isoforms whose estimated abundance falls below a minimum fraction of corresponding gene expression (default minimum of 1% presumed biologically relevant) are removed, and remaining isoforms are re-quantified. This process effectively reabsorbs read support from filtered isoforms into remaining compatible models and continues until all retained isoforms exceed the minimum gene-fractional expression threshold.

To further improve robustness under conditions of high read ambiguity, the EM-based quantification incorporates additional modeling constraints informed by long-read sequencing characteristics. These include regularization to stabilize abundance estimates when reads are compatible with multiple isoforms, as well as weighting of ambiguous read assignments based on agreement with transcript 3′ termini, reflecting the increased reliability of read evidence near polyadenylation sites based on 3’ poly-A primed cDNA synthesis. This approach prioritizes biologically informative read support while mitigating overconfidence arising from partial or weakly informative alignments.

LRAA supports three execution modes within the same algorithmic framework. In quantification-only mode, isoform discovery is bypassed and reads are assigned directly to a provided reference annotation using EM-based abundance estimation. In reference-guided mode, known transcript structures are incorporated into splice graph construction to guide discovery of additional isoforms while retaining sensitivity to novel transcript variants. In annotation-free (de novo) mode, isoform discovery and quantification rely solely on long-read alignments and the reference genome sequence, without requiring any prior transcript annotation. Across all modes, the same splice-graph representation, read compatibility modeling, and EM-based quantification strategy are used.

### Benchmarking isoform identification and quantification accuracy and study design

Benchmarking long-read isoform identification and quantification methods is challenging because no single dataset simultaneously provides complete ground truth and the complexity of real biological data. Different benchmarking substrates therefore offer complementary advantages, with each addressing distinct aspects of the problem.

Simulated long-read datasets provide complete knowledge of the isoform from which each read was generated, enabling unambiguous evaluation of isoform identification and abundance estimation even when reads are partial or compatible with multiple isoforms. However, simulated data lack many features of real sequencing experiments, including heterogeneous degradation patterns and the full complexity of transcript processing observed in biological samples, while platform-specific error profiles are necessarily modeled rather than directly observed.

Conversely, genuine long-read RNA sequencing data capture the biological and technical complexity that isoform analysis methods are designed to address, including variable transcript boundaries, partial transcripts, and technology-specific error characteristics. These datasets are therefore essential for evaluating practical performance in biologically relevant settings, but they lack definitive isoform-level ground truth. In particular, it is generally not possible to determine with certainty which isoform a partial read originated from, nor to enumerate all true full-length transcript structures present in a complex sample.

Synthetic spike-in standards, such as SIRVs and Sequins, occupy an intermediate position. These controls consist of real sequence data with defined transcript structures, enabling assessment under controlled conditions. However, because individual reads are not barcoded, the transcript of origin for each read is not directly observable, and these molecules are not expressed within native cellular contexts, limiting their ability to capture biological processing and degradation.

To bridge these gaps, we propose a new benchmarking framework based on Multiplexed Overexpression of Regulatory Factors (MORFs), which provides biologically expressed, barcoded isoforms for which the transcript of origin is known for every sequenced read. MORFs combine key advantages of simulated and real datasets by enabling unambiguous read-level truth assignment while preserving endogenous transcription, degradation patterns, and sequencing error characteristics. This framework allows direct evaluation of isoform discovery, read assignment, and abundance estimation under conditions that closely resemble real experimental data.

Across these benchmarks, definitions of true positives, false positives, and false negatives necessarily depend on both the properties of the dataset and the execution mode being evaluated. For simulated and MORF-based benchmarks, isoform– and read-level truth is explicitly defined, enabling direct assessment of identification and quantification accuracy. In contrast, for genuine whole-transcriptome datasets lacking definitive ground truth, evaluation focuses on structure-based comparisons and quantitative concordance rather than absolute correctness. For reference-guided analyses, isoforms provided as input are excluded from discovery-based performance metrics to ensure that evaluation is restricted to transcripts unknown to the method a priori. Below, we apply all these benchmarking strategies to enable evaluation of isoform identification and quantification performance across the diverse data types and analytical modes.

### Benchmarking isoform identification and quantification accuracy using simulated and genuine PacBio and ONT long isoform reads and reference data sets

To evaluate isoform identification and quantification performance across diverse experimental conditions, we benchmarked LRAA and related methods using PacBio and Oxford Nanopore Technologies (ONT) long-read reference datasets analyzed in the following progression. We first examined simulated long-read datasets, which provide complete knowledge of transcript structures and read origins, enabling direct evaluation of isoform identification and abundance estimation under controlled error models. We next assessed performance using SIRV spike-in controls, consisting of real long-read sequencing data generated from defined transcript mixtures that capture platform-specific error characteristics while retaining known reference structures. Finally, we analyzed Multiplexed Overexpression of Regulatory Factors (MORFs) datasets, which provide biologically expressed reference transcripts with read-level origin information through transcript barcoding.

### Simulated long RNA-seq at variable sequencing error

We evaluated isoform identification and quantification accuracy using simulated long-read strand-specific RNA-seq datasets, which provide a complete and unambiguous ground truth and enable controlled assessment of the impact of sequencing error and read truncation on performance. Long-read data were simulated for mouse and *Arabidopsis thaliana* transcriptomes with platform-specific 3’ and 5’ end variable truncation at error-free conditions and at increasing total error rates of 1.6%, 5.0%, and 8.5%. Across all simulations and execution modes, LRAA consistently ranked among the top-performing methods for both quantification and isoform identification (**Figure 2, Supplementary Figure 2**). In quantification-only mode, most methods performed well at low error rates but diverged as error increased, with several showing marked degradation, whereas LRAA maintained stable accuracy. For reference-guided and annotation-free isoform identification, overall accuracy declined with increasing error for all methods and as a result of read incompleteness, but LRAA showed a more balanced trade-off between recall and precision, yielding consistently strong F1 scores across error profiles. While informative for contrasting method performance on tractable inputs, simulated data are imperfect models of genuine sequencing data and do not substitute for benchmarking on real datasets.

**Figure 2:**
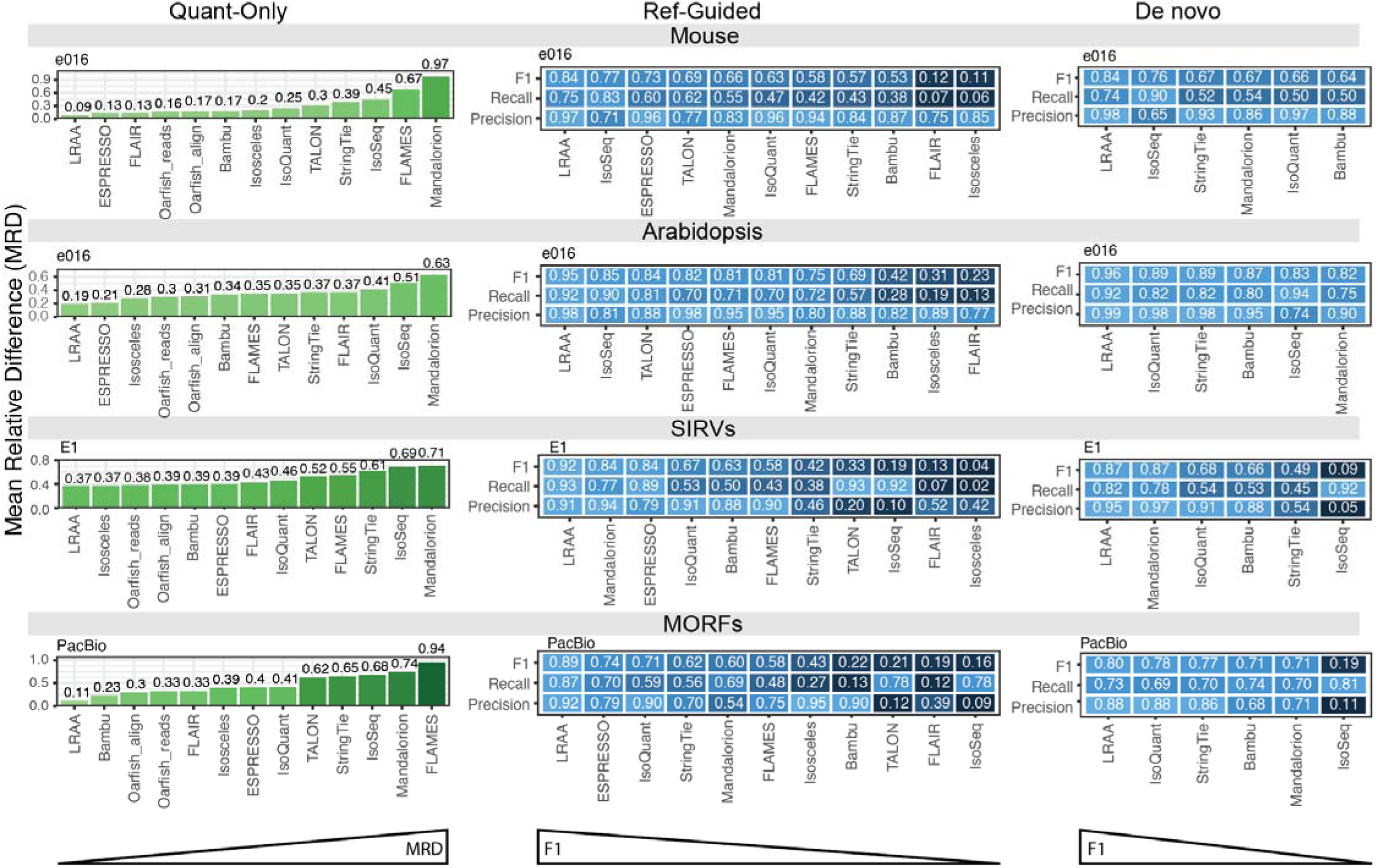
Isoform identification and quantification accuracy based on long read RNA-seq derived from simulated reads (Mouse and Arabidopsis) or genuine RNA-seq for sequencing standards. Quantification (left column), reference annotation-guided isoform identification (center column), and reference annotation-free (de novo) isoform identification (right column) accuracies are indicated for evaluated methods based on the following data sets (top-to-bottom): simulated mouse and simulated Arabidopsis PacBio reads with 1.6% error, and genuine PacBio Kinnex sequencing for SIRVs and MORFs. Methods are ranked from left to right according to top accuracy based on mean relative difference for quantification and F1 score for isoform identification.

### Spike-In RNA Variants (SIRVs)

We next evaluated isoform identification and quantification accuracy using Spike-In RNA Variants (SIRVs), which provide a defined set of transcript structures and relative abundances while retaining real sequencing and alignment characteristics. We analyzed PacBio Kinnex sequencing data for the SIRV mixes E0, E1, and E2, spanning equimolar to highly skewed expression distributions (https://www.lexogen.com/store/sirv-set1/). Across all three execution modes, quantification-only, reference-guided isoform identification, and annotation-free de novo discovery, LRAA is consistently ranked as a top-performing method (**Figure 2, Supplementary Figure 3**), outperforming other methods in both recall and precision in most settings. In quantification-only mode, LRAA achieved the highest accuracy across all SIRV mixes, with other probabilistic approaches including Oarfish, Isosceles, ESPRESSO, and Bambu also performing strongly. For isoform identification, performance varied by execution mode and expression mix. Mandalorion slightly outperformed LRAA in de novo identification for the E0 mix, and ESPRESSO showed marginally higher accuracy in the reference-guided E2 evaluation, while LRAA remained among the top-ranked methods across all conditions. Methods such as IsoSeq and TALON exhibited high sensitivity but lower precision, resulting in reduced overall accuracy. While SIRVs provide a useful intermediate benchmark, they remain synthetic constructs and do not fully reflect native cellular processing of transcripts, which motivated us to explore MORFs as benchmarks described below.

### MORFs

To benchmark isoform identification and quantification using genuine biological transcripts with unambiguous ground truth, we leveraged the Multiplexed Overexpression of Regulatory Factors (MORF) library (Joung et al., 2023). This resource comprises 3,548 human transcription factor isoforms representing 1,836 genes, expressed as lentiviral clones and each containing a unique isoform-specific barcode embedded at the 3′ end of the transcript (**Supplementary Figure 4a**). These barcodes enable definitive assignment of every sequenced read, including partial reads, to its isoform of origin, thereby overcoming a central limitation of synthetic spike-in controls and genuine transcriptome data. We sequenced the MORF library after expression in HEK293 cells using both PacBio Kinnex and Oxford Nanopore platforms and evaluated isoform identification and quantification accuracy across quantification-only, reference-guided, and annotation-free de novo execution modes (**Figure 2, Supplementary Figure 4b**). Across all evaluations, LRAA achieved the highest overall accuracy for both isoform identification and quantification. In the reference-guided setting for isoform discovery, LRAA was the top performer, significantly outranking all other methods, with ESPRESSO being the next most highly ranked. In contrast, for *de novo* mode, IsoQuant and StringTie proved to be the strongest alternatives. For quantification, Bambu and Oarfish ranked next best after LRAA. As observed for SIRVs, IsoSeq and TALON exhibited high recall but substantially lower precision, resulting in reduced overall accuracy. Overall, MORFs provided a more biologically informed benchmark that revealed shifts in relative method performance compared with SIRVs, most notably for Mandalorion, while also confirming that a subset of methods, including LRAA, consistently ranked among the top performers.

### Whole transcriptome sequencing of cell lines and reference standards

While simulations, SIRVs, and MORFs enable benchmarking against defined truth sets, the primary use case for long-read isoform analysis methods is whole-transcriptome sequencing, where ground truth isoform structures and expression levels are not known. To evaluate performance in this etting, we analyzed bulk long-read transcriptomes generated using PacBio Kinnex for the BT474 and K562 cell lines, the lymphoblastoid sample GM24385 provided as Genome in a Bottle RNA reference sample for individual HG002 (https://www.nist.gov/programs-projects/genome-bottle), the Universal Human RNA Reference (UHRR) (Xu et al., 2014), and Oxford Nanopore strand-specific cDNA sequencing for the A549 and MCF7 cell lines from the SG-NEx project (Chen et al., 2025). Isoform identification accuracy was assessed using both reference-guided and annotation-free *de novo* execution modes, with proxy truth sets defined based on agreement across methods and known reference annotations (see **Methods**). Across all PacBio Kinnex datasets, LRAA ranked as the top-performing method for isoform identification in both execution modes, while for the ONT A549 dataset StringTie marginally outperformed LRAA, with comparable performance observed for MCF7 (**Figure 3**). Because no absolute expression ground truth is available for these datasets, we evaluated quantification consistency by comparing isoform-level expression profiles across methods using Pearson correlation. LRAA-derived expression profiles clustered closely with those from methods that performed best in quantification benchmarks with defined truth sets, including Oarfish, Isosceles, and Bambu. Overall, these results demonstrate that LRAA performs robustly across whole-transcriptome datasets and sequencing platforms, positioning LRAA as a highly effective approach for long-read transcriptome analysis.

**Figure 3:**
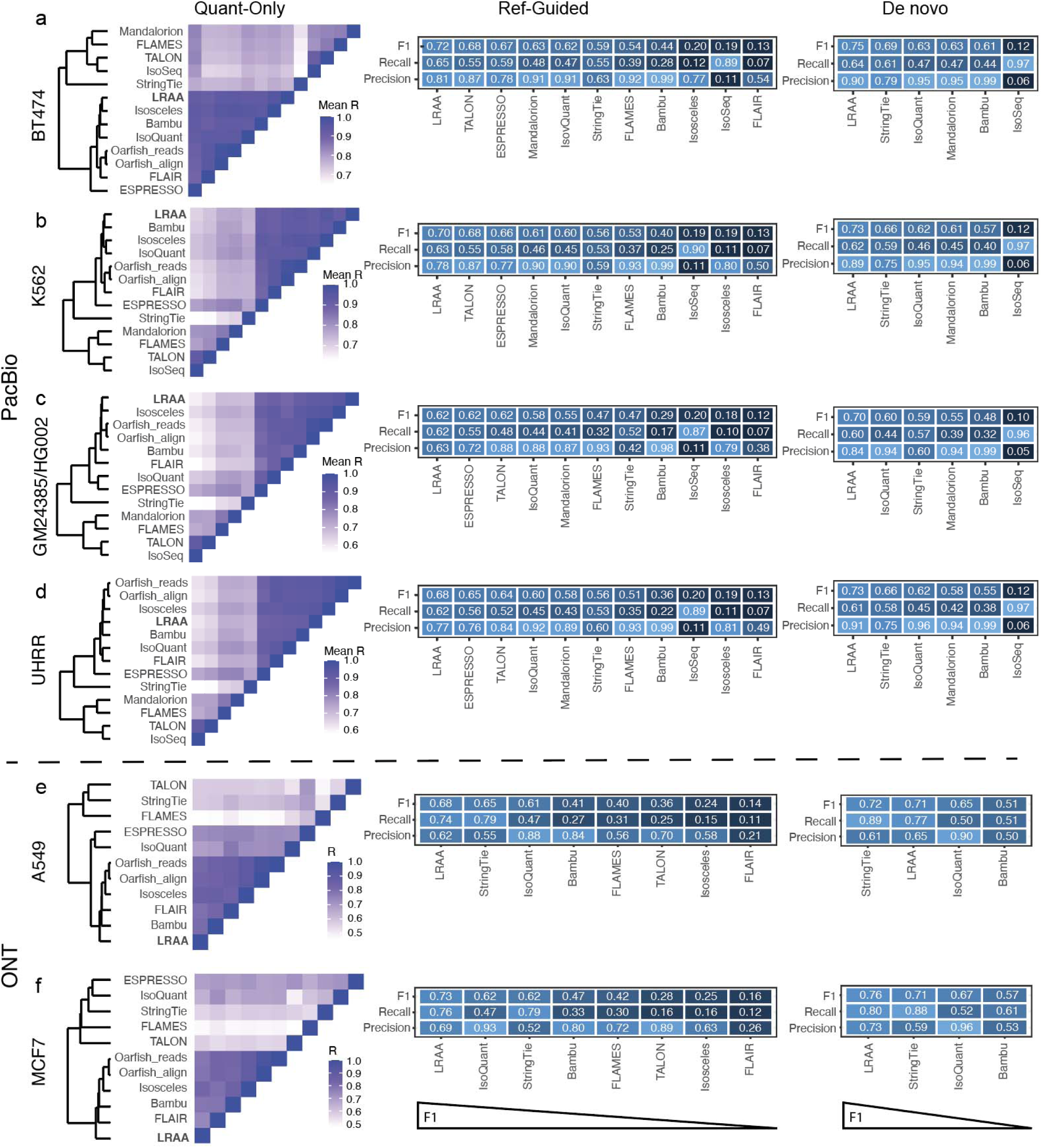
Benchmarking isoform identification and quantification accuracy using whole transcriptome sequencing of human cell lines and RNA reference samples. Transcript quantification (left column), reference annotation-guided (center column) and reference annotation-free (de novo) isoform identification (right column) were evaluated for long read isoform analysis methods based on whole transcriptomes sequenced using PacBio Kinnex (top section) or ONT (bottom section). PacBio Kinnex sequenced transcriptomes correspond to (top-to-bottom) cell lines (a) BT474 and (b) K562 and (c) reference standards Genome In a Bottle RNA sample for GM24385/HG002 and (d) commercially available Universal Human RNA Reference (UHRR) from Thermo Fisher. ONT sequenced cell lines correspond to (top-to-bottom) (e) A549 and (f) MCF7. Expression quantification was evaluated by computing the Pearson correlations for isoform expression (log(TPM+1) values) and clustered by similarity. Isoform identification accuracy was estimated using proxy truth sets based on shared vs. uniquely identified isoforms (see **Methods**).

### Application of LRAA for de novo or reference-guided isoform identification in single-cell and single-nucleus long-read whole transcriptome sequencing

Application of long-read transcriptome sequencing at single-cell and single-nucleus resolution enables isoform-level analysis in heterogeneous tissues but introduces additional computational challenges relative to bulk data (Al’Khafaji et al., 2024; P. Gupta et al., 2024; Hardwick et al., 2022; Kumari et al., 2024). In particular, individual cells contribute limited read depth, rare isoforms may be confined to specific cell populations, and single-nucleus data are enriched for partially processed transcripts. To address these challenges, we developed an LRAA single-cell workflow that integrates isoform identification and quantification with cell clustering, allowing isoform discovery to be informed by cell-type-specific expression patterns (**Figure 4**).

**Figure 4.**
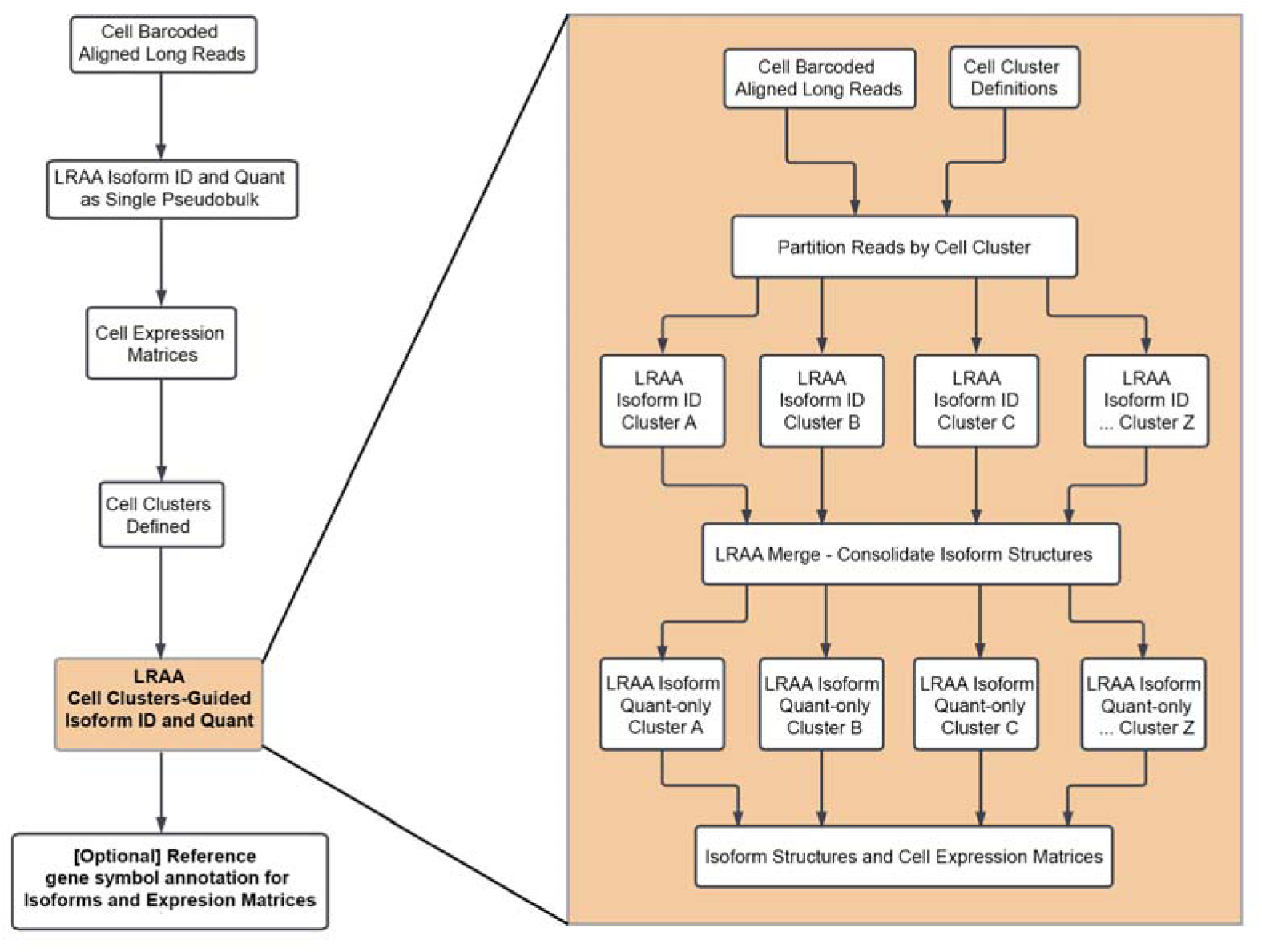
Pipeline for application of LRAA to single-cell transcriptomics data for novel isoform discovery. LRAA is first leveraged for isoform identification and quantification in pseudobulk mode (yellow). Cell barcodes derived from isoform-assigned reads are leveraged to define cell clusters. LRAA is then used on a second phase where isoform identification is performed separately on each cell cluster, using the isoform structures defined in the initial phase for reference-guided isoform identification. The cell cluster-based isoforms are subsequently merged and unified followed by a final round of cell cluster-based quantification to assign reads (and cell barcodes) to the unified isoform set, ultimately retaining cell-type specific isoform structures.

The LRAA single-cell workflow operates in two phases. In the first phase, LRAA is applied in pseudobulk mode to perform isoform identification and quantification using either reference-guided or annotation-free execution modes. Gene-level expression estimates derived from this phase are then used to define cell clusters based on cell-barcoded read assignments. In the second phase, isoform identification is performed separately within each cell cluster, enabling recovery of isoforms that are low abundance in the full pseudobulk but enriched within specific cell populations. Isoforms identified across clusters are subsequently merged into a unified isoform set, followed by a final round of cluster-aware quantification to generate gene-, isoform-, and splice-pattern-level expression matrices for downstream analysis.

We applied this workflow to two distinct long-read single-cell datasets generated using PacBio Kinnex sequencing of 10x 3′ libraries: peripheral blood mononuclear cells (PBMCs), representing a well-characterized and transcriptionally diverse cell population, and single nuclei isolated from frontal cortex tissue of an FTD donor, representing a challenging setting dominated by intronic reads and incomplete transcript processing.

### Isoform Detection via Single-Cell Kinnex from PBMCs

We first applied the LRAA single-cell workflow to a PBMC sc-Kinnex dataset generated using PacBio sequencing of a 10x 3′ library totaling 82M reads for 10k cells. We examined the characteristics of individual long-read alignments relative to the reference annotation using an LRAA SQANTI-like classification (see **Methods**). A substantial fraction of reads corresponded to fully spliced matches (FSM, 29%) and incompletely spliced matches (ISM, 9%), with additional reads mapping to intronic or antisense regions (**Figure 5a**).

**Figure 5:**
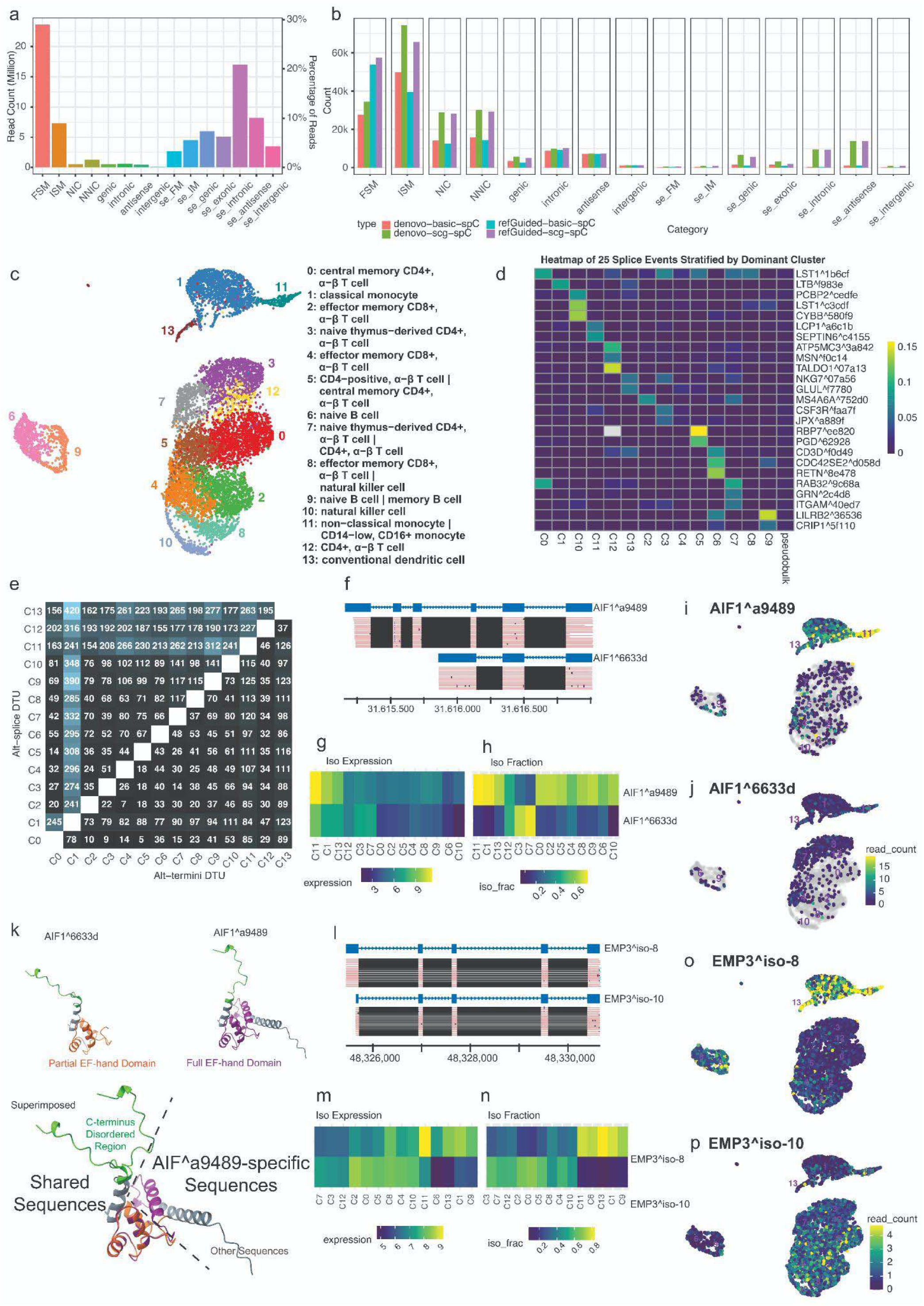
Isoform identification and quantification in PBMCs via LRAA with Kinnex scRNA-Seq. (a) Counts and percentages of Kinnex reads assigned to gene structures or genome regions according to LRAA SQANTI-like categories. (b) Counts of LRAA-reconstructed isoforms from reference annotation-free (de novo) or reference annotation-guided workflows assigned to SQANTI-like categories. Isoforms with identical splice patterns but different termini boundaries are counted non-redundantly as collapsed. (c) UMAP for PBMCs based on LRAA-derived gene expression in quantification-only mode with dominant inferred cell types indicated. (d) Examples of isoforms estimated to have <1% isoform fraction in pseudobulk mode but >10% isoform fraction in at least one cell cluster context. (e) Counts of genes with detected differential transcript usage (DTU, upper triangle) or having identical splicing patterns but alternative termini usage (ATU, lower triangle) by cell cluster comparison. (f-j) DTU detected for two isoforms of the AIF1 gene with (f) isoform structures and full-length Kinnex reads, (g) isoform expression, (h) isoform fraction, and (i,j) cellular expression shown for each isoform. (k) Structural modeling of AIF1 proteoforms. (l-p) ATU in the form of alternative transcriptional start sites detected for two isoforms of the EMP3 gene with (l) isoform structures and full-length Kinnex reads, (m) isoform expression, (n) isoform fraction, and (o,p) cellular expression shown for each isoform.

LRAA was run in both annotation-free de novo and reference-guided modes. To ensure consistent downstream comparisons, cell clustering was derived from an initial quantification-only run, and the same cluster assignments were used for subsequent cluster-guided isoform identification and quantification in both execution modes (**Figure 5c**). We compared isoform structures obtained from the initial pseudobulk phase and the final cluster-guided phase of the workflow. The cluster-guided phase yielded modest increases in FSM isoforms (7%) but substantially increased the number of ISM (66%) and novel isoforms (114%), while also contributing many more single-exon transcripts, mostly which were localized within introns or oriented antisense to annotated transcripts (**Figure 5b**).

Notably, the cluster-guided phase enabled recovery of isoforms that were present at low fractional abundance in the pseudobulk analysis but became prominent within specific cell clusters. Many such isoforms failed to meet minimum isoform fraction thresholds in the initial phase but exceeded these thresholds when evaluated within cluster-specific contexts (**Figure 5d**). Reference-guided execution increased the number of FSM isoforms relative to annotation-free discovery but had limited impact on the recovery of ISM and novel isoforms.

Using the reference-guided, cluster-guided isoform set, we examined differential transcript usage across PBMC cell clusters. We identified genes exhibiting shifts in dominant isoform usage driven by alternative splicing (mean 154 genes per pair cell clusters), as well as genes with identical splice patterns but alternative transcription start or end site usage (mean 60 genes per pair cell clusters) (**Figure 5e**). Representative examples include isoform switching of *AIF1* between non-classical monocytes and subsets of T cells, and alternative transcription start site usage of *EMP3* across monocytes, dendritic cells, B cells, and T cell populations (**Figure 5f–p**).

To further characterize the functional implications of alternative isoform usage, we examined the protein structural differences between the two proteoforms encoded by alternatively used *AIF1* isoforms across PBMC cell types (**Figure 5k**). Structural modeling revealed that AIF1^6633d encodes a truncated protein containing only a partial EF-hand calcium-binding domain, while AIF1^a9489 encodes the full-length protein with a complete EF-hand domain (**Figure 5k**). Superimposition of the two structures showed extensive overlap, with AIF1^a9489 containing additional structured and unstructured regions that may confer distinct calcium-binding properties and protein-protein interaction capabilities. Both proteoforms matched known reference annotations in GENCODE v47, demonstrating that reference-annotated alternative splicing can generate proteoforms with substantially different structural and potentially functional properties.

These findings motivated us to systematically investigate the landscape of novel proteoforms, those arising from unannotated splice junctions or intron retention events, that may expand proteomic diversity beyond the reference annotation. To identify novel proteoforms in human PBMCs, we applied our classification pipeline to complete ORF protein sequences at three length thresholds (≥50 AA, ≥100 AA, ≥200 AA). At the ≥50 AA threshold, 74,212 unique proteins were detected, with novel proteoforms representing 12.6% (9,377) of the total proteome (**Supplementary Figure 6, top**). Notably, as stringency increased to ≥100 AA and ≥200 AA, the proportion of novel proteoforms remained more or less consistent at 13.2% (6,927 of 52,490) and 11.9% (3,583 of 30,147), respectively, indicating that novel splicing events generate proteoforms across the full spectrum of protein sizes. Analysis of novel proteoform composition revealed that novel splice junctions accounted for 81.5-84.9% of novel proteoforms across all thresholds, while intron retention events contributed 14.5-17.9% (**Supplementary Figure 6, bottom**).

### Isoform Detection in Single-Cell Nuclei from an FTD Brain Frontal Cortex Sample

We next applied the LRAA single-cell workflow to long-read single-nucleus RNA-seq data generated from frontal cortex tissue of an FTD donor. Single-nucleus sequencing enables transcriptional profiling of cell types that are difficult to capture using whole-cell approaches, but the resulting data are enriched for unspliced and partially processed transcripts, posing challenges for isoform-level analysis.

The FTD frontal cortex dataset was generated using PacBio Kinnex sequencing of a 10x 3′ single-nucleus library, yielding approximately 55 million long reads for 6.5k nuclei. In contrast to the PBMC sc-Kinnex data, the majority of reads were classified as intronic (42%) according to LRAA SQANTI-like read classification, with substantially fewer reads corresponding to fully spliced (3%) or incompletely spliced matches (4%) (**Figure 6a**). Despite this, LRAA identified large numbers of isoforms in both annotation-free and reference-guided execution modes, with the reference-guided analysis yielding ∼3X as many fully spliced isoforms (**Figure 6b**).

**Figure 6.**
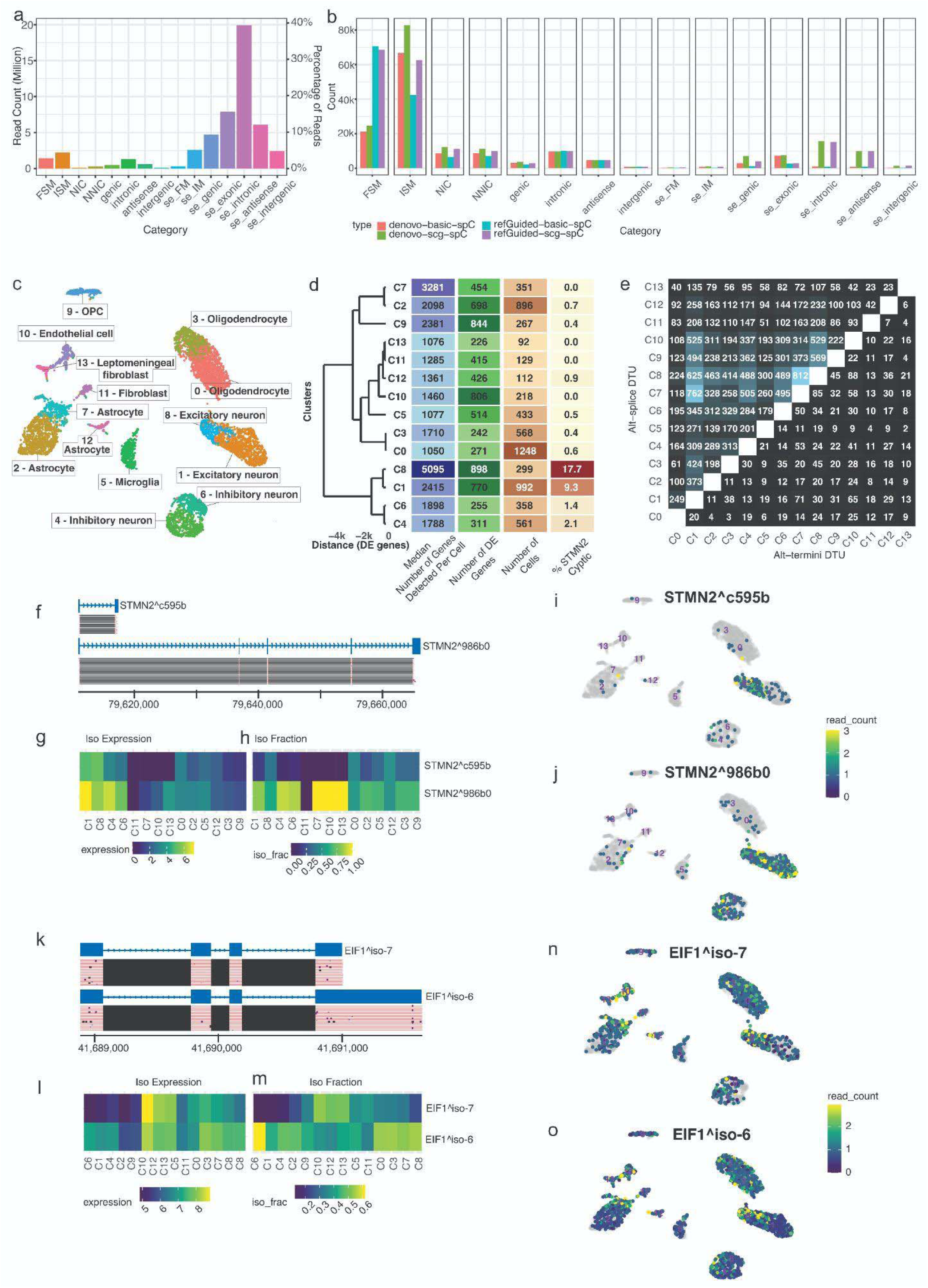
Isoform identification and quantification in an FTD brain frontal cortex via LRAA with Kinnex snRNA-Seq. (a) Counts and percentages of Kinnex reads assigned to gene structures or genome regions according to LRAA SQANTI-like categories. (b) Counts of LRAA-reconstructed isoforms from reference annotation-free (de novo) or reference annotation-guided workflows assigned to SQANTI-like categories. (c) UMAP for FTD frontal cortex sample based on LRAA-derived gene expression in quantification-only mode with dominant inferred cell types indicated. (d) Overview of cell cluster properties and dendrogram showing relationships between cell clusters based on numbers of differentially expressed genes defined as cluster-specific biomarkers (p_val_adj < 1e-5 & abs(avg_log2FC >= 2)). (e) Counts of genes with detected differential transcript usage (DTU, upper triangle) or having identical splicing patterns but alternative termini usage (ATU, lower triangle) by cell nuclei cluster comparison. (f-j) DTU detected for two isoforms of the *STMN2* gene with (f) isoform structures and full-length Kinnex reads, with the pathogenic cryptic isoform at top and canonical *STMN2* isoform at bottom, (g) isoform expression, (h) isoform fraction, and (i,j) cellular expression shown for each isoform. (k-o) ATU in the form of alternative polyadenylation sites detected for two isoforms of the *EIF1* gene with (k) isoform structures and full-length Kinnex reads, (l) isoform expression, (m) isoform fraction, and (n,o) cellular nuclei expression shown for each isoform.

Comparison of the initial pseudobulk phase and the cluster-guided phase showed limited differences in the number of fully spliced isoforms, but substantial increases in incompletely spliced (24% increase for de novo and 48% increase for reference-guided) and novel isoforms (36% increase for de novo and 54% increase for reference-guided) following cluster-guided isoform discovery (**Figure 6b**). The abundance of intronic reads did not lead to a proportional increase in intronic isoform calls, consistent with filtering based on read support, transcription start site evidence, and transcript end site evidence.

Gene-level expression profiles derived from the quantification-only phase were used to identify 14 distinct cell clusters corresponding to major brain cell types, including excitatory and inhibitory neurons, astrocytes, oligodendrocytes, microglia, endothelial cells, and fibroblasts (**Figure 6c,d**). Several of these cell types were further resolved into transcriptionally distinct subpopulations, often differing substantially in the number of expressed genes per nucleus (**Figure 6d**). Pairwise differential expression analysis between paired cell type subpopulations coupled with functional enrichment analysis indicated one subpopulation to be generally more active than the other coupled with stress responses. We refer to these as active or quiescent subpopulations below.

Using the reference-guided, cluster-guided isoform set, we examined differential transcript usage across cell clusters. Despite the predominance of intronic reads, we identified genes exhibiting significant shifts in isoform usage between cell types (**Figure 6e**). Among these, we detected the pathogenic cryptically spliced isoform of *STMN2*, a hallmark of TDP-43 proteinopathy in FTD (**Figure 6f–j**) (Klim et al., 2019; Melamed et al., 2019; Prudencio et al., 2020). Expression of *STMN2* transcripts was most prevalent in excitatory neuron clusters 8 and 1, where *STMN2* expression was detected in 42% and 38% of nuclei, respectively. However, the fraction of nuclei expressing the cryptically spliced isoform differed substantially between these clusters, with 18% of active cluster 8 nuclei showing evidence of the cryptic isoform compared to 9% in quiescent cluster 1. Inhibitory neuron clusters (active) 4 and (quiescent) 6 also exhibited *STMN2* expression in 19% and 11% of nuclei, respectively, but the cryptically spliced isoform was detected in only 2% and 1% of nuclei in these clusters.

In addition to alternative splice pattern usage, we identified cases of alternative transcript end site usage without changes in splice structure. One such example involved *EIF1*, where isoforms with identical splicing patterns but distinct polyadenylation sites were detected across multiple cell types, with the most pronounced shifts observed between oligodendrocytes and endothelial cells (**Figure 6k–o**).

We applied the same proteoform analysis pipeline to FTD brain frontal cortex samples. Novel proteoforms represented 5.2% (3,283 of 63,153), 5.7% (2,408 of 42,560), and 4.7% (1,126 of 23,844) of the total proteome at ≥50 AA, ≥100 AA, and ≥200 AA thresholds, respectively (**Supplementary Figure 7**). Consistent with PBMC samples, novel junctions predominated across all thresholds (66.6-74.1%), while intron retention contributed 25.2-32.9%.

## Discussion

Long-read transcriptome sequencing has transformed transcriptome analysis by enabling direct observation of full-length isoforms rather than inference from short-read fragments. Recent gains in throughput and accuracy have extended these capabilities to single-cell and single-nucleus settings, creating new opportunities to study isoform regulation in heterogeneous tissues and disease. At the same time, these advances have exposed persistent computational challenges, including partial transcript coverage, ambiguous read-to-isoform compatibility, and dependence on incomplete reference annotations. Addressing these challenges requires methods that integrate accurate isoform discovery with principled quantification and that generalize across sequencing platforms and experimental contexts.

Here we present LRAA as a unified framework for isoform identification and quantification from long-read RNA sequencing data spanning bulk, single-cell, and single-nucleus applications. LRAA integrates splice-graph–based structural modeling with an expectation–maximization framework for abundance estimation, enabling probabilistic resolution of ambiguous reads while retaining sensitivity to novel isoforms. Across a broad range of benchmarking datasets, LRAA demonstrated robust performance across execution modes, sequencing technologies, and biological settings. In evaluations that included leading methods assessed by the LRGASP Consortium, such as IsoQuant, Bambu and StringTie2, LRAA ranked among the strongest-performing approaches in quantification-only, reference-guided, and de novo analyses. In cases where LRAA did not achieve the highest overall accuracy, its performance closely matched that of the top-ranked method.

Rigorous benchmarking of long-read isoform analysis methods remains challenging because no single dataset simultaneously provides complete ground truth and biological realism. Simulated datasets offer unambiguous isoform-of-origin information and are valuable for controlled evaluation of algorithmic behavior, including sensitivity to sequencing error. However, simulations necessarily lack native transcriptional processing and degradation. Synthetic spike-in controls such as SIRVs incorporate real sequencing characteristics and defined expression ratios, but remain artificial constructs that do not fully reflect cellular transcript processing. As a result, method performance on simulations or spike-ins does not always generalize to biologically complex datasets To address this gap, we leveraged the MORF library as a biologically informed benchmarking substrate. MORFs consist of thousands of human isoforms expressed in living cells, each tagged with an isoform-specific barcode that enables unambiguous assignment of every read, including partial reads, to its isoform of origin. This design combines the ground-truth clarity of simulations with the biological context of native transcription. MORF benchmarking revealed certain tools to have notable differences in relative method performance compared with benchmarks on SIRVs, while others, such as LRAA, maintained consistently strong performance across benchmarking conditions. These results underscore the value of biologically inspired benchmarks for realistic evaluation of long-read isoform analysis tools.

Beyond controlled benchmarks, we evaluated LRAA on whole-transcriptome datasets from multiple human cell lines and reference RNA samples sequenced using both PacBio and Oxford Nanopore technologies. In this setting, where absolute ground truth is unavailable, LRAA showed strong performance for isoform identification and produced expression profiles that closely matched those of methods previously shown to perform well under benchmarks with defined truth sets. These results indicate that LRAA generalizes effectively to real-world transcriptome analyses and performs robustly across sequencing platforms without requiring platform-specific workflows.

A major advance of this work is the extension of isoform-resolved long-read analysis to single-cell and single-nucleus transcriptomics. Single-cell long-read data are sparse, and biologically relevant isoforms may be confined to specific cell populations and obscured in pseudobulk analyses. By integrating isoform discovery with cell clustering, LRAA enables recovery of isoforms that are low abundance at the sample level but prominent within particular cell types. In PBMCs, this approach revealed cell-type-specific isoform usage driven by both alternative splicing and alternative transcript start or end site selection.

Single-nucleus transcriptomics presents additional challenges, as nuclear RNA is enriched for unspliced and partially processed transcripts. Despite these constraints, LRAA identified biologically meaningful isoform usage in single-nucleus data from FTD frontal cortex tissue. In particular, we detected the pathogenic cryptically spliced isoform of *STMN2*, a hallmark of TDP-43 proteinopathy, with marked differences in prevalence across neuronal subpopulations. These observations demonstrate that long-read single-nucleus sequencing, coupled with appropriate computational modeling, can resolve disease-relevant isoform dysregulation in tissues that are poorly represented in conventional single-cell approaches.

Our identification of novel proteoforms, protein isoforms arising from unannotated splice junctions or intron retention events, revealed that ∼13% of the PBMC proteome and ∼5% of the FTD frontal cortex tissue proteome may consist of sequences not present in current reference annotations. These novel proteoforms represent complete ORFs with canonical start and stop codons, suggesting translational competence, though translatomics and proteomics validation will be essential to confirm bona fide expressed proteins and to establish their functional relevance in health and disease.

Several limitations remain. Although LRAA models alternative transcription start and end sites, benchmarking here focused primarily on splice-pattern accuracy, reflecting ongoing challenges in reliably assessing transcript termini in the presence of RNA degradation and protocol-specific biases. Single-exon isoforms remain difficult to evaluate in de novo discovery settings, necessitating conservative filtering to limit false positives. Continued improvements in sequencing protocols, library preparation, and benchmarking standards will further advance isoform-resolved transcriptomics.

As long-read sequencing is increasingly integrated with single-cell, single-nucleus, and emerging spatial technologies, the ability to accurately resolve isoform structure and usage at scale will become central to transcriptome analysis. By combining robust isoform discovery, principled quantification, and applicability across experimental contexts, LRAA represents a foundational framework for dissecting isoform regulation across development and disease.

## Methods

### LongReadAlignmentAssembler (LRAA) algorithm

We developed LongReadAlignmentAssembler (LRAA) as a method for transcript isoform discovery and quantification from long-read RNA sequencing data. LRAA reconstructs full-length transcript isoforms by constructing a splice graph from long RNA-seq read alignment-supported exons and introns, enumerating read-consistent paths through this graph, and selecting high-confidence isoforms via iterative best-path extraction. Isoform abundances are estimated using an expectation-maximization (EM) algorithm that resolves ambiguous read assignments.

LRAA has three primary execution modes: (1) annotation-free isoform discovery and quantification, requiring only aligned reads and a reference genome; (2) reference annotation-guided isoform discovery and quantification, where known gene models inform splice-graph construction and filtering; and (3) quantification-only mode, where isoform discovery is bypassed and reads are directly assigned to reference isoforms for abundance estimation via EM.

LRAA was primarily developed for PacBio HiFi data, which provides high base accuracy (>99%) and enables precise inference of transcription start sites (TSS) and polyadenylation (PolyA) sites from read termini. The method operates in HiFi mode (the ––HiFi flag) for PacBio datasets, which activates stringent filtering parameters optimized for high-accuracy reads. LRAA also supports Oxford Nanopore Technologies (ONT) data in default mode, which relaxes filtering thresholds to accommodate higher sequencing error rates. Unless otherwise noted, parameter values and algorithmic decisions described below reflect the HiFi mode configuration, and thresholds defined below are user-configurable.

### Input and preprocessing

LRAA requires coordinate-sorted, indexed BAM files of long RNA-seq reads aligned to a reference genome (typically via minimap2), along with the reference genome in FASTA format. Reference gene annotations in GTF format are required for quantification-only mode or reference annotation-guided isoform discovery. Annotation-free isoform identification and quantification requires only the input alignment BAM and reference genome files.

### Filtering of alignments

Stringent quality filters are applied during read ingestion. Reads below the minimum mapping quality of 1 are discarded. In HiFi mode, reads below 98% identity are filtered out to leverage high base accuracy and reduce alignment artifacts; for ONT data, this identity threshold is relaxed to 80%. Secondary, duplicate, and QC-failed alignments are excluded.

### Normalization of read coverage for isoform discovery

To reduce computational requirements in ultra-high-coverage regions, alignments are down-sampled to a maximum per-base coverage of 1000× using a strand-aware normalization procedure applied independently per strand before splice-graph construction. The original, unnormalized BAM is used for isoform abundance estimation, ensuring that quantification accuracy is preserved.

### Correction of alignments at soft-clipped termini

Long reads often contain soft-clipped bases at alignment boundaries when short segments fail to align across splice junctions to adjacent exons. Optionally, LRAA examines soft-clipped termini in the splice graph context and realigns them to connected exon segments to incorporate legitimate exonic extensions. Soft-clipped regions at 3′ ends containing ≥7 adenines (or thymines for the reverse strand), comprising ≥80% of the sequence, are identified as post-transcriptional PolyA tails and removed, revealing the genomic polyadenylation site.

### Splice-graph construction

For each genomic contig and strand, LRAA constructs a directed acyclic graph representing the exon–intron architecture inferred from read alignments. Graph nodes correspond to contiguous genomic intervals supported by read coverage (exon segments) or splice junctions (introns). Adjacent exon segments separated by less than 50 bp with no intervening splicing are merged. Introns supported by spliced read alignments become nodes that connect the corresponding exon nodes in the graph.

Read support is quantified differently for exon and intron nodes. For introns, each read spanning the junction increments the intron’s support count. For exons, genomic base coverage is accumulated across all aligned reads; the mean coverage across the exon segment’s coordinates is then computed and assigned as the node’s support value. These support metrics are used throughout downstream filtering and scoring steps.

In HiFi mode, TSS and PolyA sites are explicitly inferred from read boundary clustering. Reads terminating with minimal or no soft-clipping (default 0 bases soft-clipped) within a 50 bp window are aggregated to define boundary candidates. A candidate site is retained if supported by at least 5 reads representing ≥5% of gene-level read coverage. TSS and PolyA nodes are integrated as first-class graph features that influence downstream isoform selection. For ONT data, TSS and PolyA inference is disabled; terminal boundaries are instead refined empirically after reconstruction.

To remove spurious features arising from alignment errors, the splice graph is pruned and filtered using several criteria. Introns supported by fewer than 2 reads are removed unless they match known junctions from a provided reference annotation. For competing splice junctions at the same donor or acceptor site, minor junctions representing less than 1% (HiFi mode) or 3% (ONT mode) of total junction support are pruned. Short terminal exon segments (≤13 bp in HiFi mode, ≤20 bp for ONT) that are not anchored by TSS or PolyA nodes are removed. Unspliced intervals bridging annotated introns are retained only if supported by ≥1% of junction coverage, distinguishing genuine intron retention from unprocessed pre-mRNA. When a reference GTF is provided, known exons and introns are integrated to further guide graph construction.

After pruning and filtering, the refined splice graph is partitioned into weakly connected components, each ideally representing a candidate gene locus. In HiFi mode, TSS and PolyA sites undergo additional filtering within each component. Sites representing less than 5% of total boundary read support within their component are removed as likely noise. TSS sites are further examined for potential RNA degradation artifacts. The graph is traversed along linear exon paths from each high-support TSS, and alternative TSS sites along this path with ≤20% of the dominant TSS support are removed as probable 5′ degradation products. After this boundary refinement, components are rediscovered to account for any resulting graph topology changes.

### Read compatibility class (RCC) definitions and RCC graph construction

Each read alignment is represented as a labeled splice graph read path, defined as an ordered sequence of splice-graph nodes encoding the exon–intron structure traversed by the read. Reads with identical splice structure are aggregated as **Read Compatibility Classes** (**RCCs**), and their labeled read paths are compiled into a directed acyclic graph (the **RCC Graph**). Each node in the RCC Graph represents a unique consecutive sequence of splice-graph nodes and is annotated with genomic coordinates, read support count, and boundary flags indicating whether the path begins or ends at an inferred TSS or PolyA site. Compatible containment relationships are established between RCCs where one path is fully encompassed within another and have identical overlapping splicing events, connecting them by an edge in the RCC graph.

Importantly, even when splice junctions are compatible and one RCC path is fully contained within another, differing TSS or PolyA boundary annotations establish incompatibilities. Compatible and contained RCC paths lacking boundary annotations, or sharing identical TSS or PolyA boundaries with their contained RCC path, are evaluated for collapsing into single RCCs based on expression thresholds, as they likely represent incomplete or degraded transcripts rather than biologically distinct isoforms.

The RCC Graph is partitioned into weakly connected components using shared splice-graph nodes. Each component represents a set of transcripts sharing at least one exon or intron and is processed independently for isoform reconstruction. To prevent combinatorial explosion in highly complex loci, components exceeding 1000 RCCs (default) are reduced to the 1000 most highly supported RCCs based on corresponding RCC read support, ensuring computational tractability while retaining the most confidently supported transcript structures.

### Isoform reconstruction

LRAA reconstructs transcript isoforms from the RCC Graph using trellis-based dynamic programming with iterative best-path extraction. The RCC Graph is unrolled into a trellis indexed by topological order, where each layer corresponds to a labeled RCC node, with top-level RCC nodes reflecting full-length or most complete isoform structures. Dynamic programming aggregates path scores across all possible routes from source to sink nodes.

Path scores integrate multiple factors: read support (primary weight), boundary consistency (bonus scores for paths with explicit TSS and PolyA boundaries), and transition penalties for weak edges or incompatible boundaries. This scoring function prioritizes high-coverage, boundary-anchored paths while permitting discovery of lower-abundance isoforms in subsequent iterations.

The highest-scoring path is identified via backtracking through the trellis and materialized as a transcript. Reads strongly explained by this transcript (≥90% compatibility) are down-weighted or removed from labeled read path counts. The trellis is rebuilt with updated counts, and the process repeats until no remaining path exceeds minimum support (default: 1 read) or until all reads are assigned.

### Isoform filtering

Reconstructed isoforms are filtered to remove very lowly expressed and low-confidence transcripts. Isoforms representing less than 1% of gene-level reads are removed unless they match annotated transcripts (reference-guided mode) with non-zero expression. This filter is applied iteratively within each gene. Read assignments and isoform fractions are recalculated via EM after each filtering round, and isoforms are sorted by expression. When a gene has more than 10 isoforms (default threshold for aggressive filtering), multiple low-abundance isoforms may be removed in a single round to accelerate convergence. Otherwise, only the lowest-expressing isoform failing the threshold is removed per round. Iteration continues until all remaining isoforms satisfy the minimum expression criteria or until no further isoforms can be removed.

Single-exon transcripts are subject to a minimum expression threshold (default: 1 TPM) to mitigate false positives from transcriptional noise. In HiFi mode, single-exon transcripts additionally require TSS or PolyA annotation, as boundary evidence available from HiFi reads is useful for distinguishing genuine monoexonic genes from transcriptional noise or alignment artifacts.

When one isoform is fully contained within another (identical splicing but shorter terminal exons), collapsing decisions depend on boundary annotations and relative expression. Contained isoforms lacking both TSS and PolyA annotations are automatically removed as likely degradation products. Contained isoforms lacking TSS but sharing the same PolyA site with their containing isoform are removed as redundant 5′ truncations. For contained isoforms with distinct boundary annotations, those representing less than 20% of the combined expression of both isoforms are removed as probable incomplete transcripts rather than genuine alternative termini. Importantly, contained isoforms with distinct TSS or PolyA boundaries that exceed this expression threshold are retained as separate isoforms. This enables discovery of biologically meaningful alternative transcription start sites and polyadenylation sites that produce overlapping but functionally distinct transcript variants.

Isoforms with 3′ termini within 10 bp of genomic A-rich sequences (≥7 consecutive adenines) are flagged as potentially internally primed, representing artifacts from oligo-dT priming within transcripts rather than at genuine PolyA tails. Flagged monoexonic isoforms are removed unless they match known 3′ ends from a provided reference annotation.

Novel isoforms (those not matching reference annotation) must be supported by at least 2 uniquely assigned reads.

### Isoform terminal exon boundary refinement

For isoforms lacking explicit TSS or PolyA annotation, corresponding terminal exon boundaries are refined using the empirical distribution of read end positions within terminal exons of supported isoforms. Terminal exon coordinates are adjusted using the following percentiles: the 10th percentile of read positions for the left boundary and the 90th percentile for the right boundary. A minimum of 7 reads must terminate in a terminal exon for percentile-based adjustment to proceed. If insufficient reads are available, the method uses the extreme (minimum/maximum) observed positions of isoform-supported reads.

### Grouping isoforms to gene loci

Reconstructed isoforms are assigned to gene loci using a two-stage clustering approach. First, isoforms are grouped by genomic overlap: transcripts sharing exonic overlap (≥1 bp) are merged into preliminary gene components. Within each initial overlap component, Leiden community detection is applied to further refine gene assignments. A binary graph is constructed in which transcripts are nodes and edges connect transcript pairs satisfying the overlap thresholds: overlap length must be ≥50% of the shorter transcript and ≥20% of the longer transcript. Leiden clustering is performed with a resolution parameter of 0.2.

For extremely large components (>1500 transcripts), Leiden is bypassed and a computationally efficient union-find algorithm is applied instead to define connected components based on the minimum isoform pairwise overlap criteria. This two-stage approach partitions the overlap graph into tightly connected subgraphs (communities) that ideally represent distinct genes whose isoforms share overlapping exons. Transcripts within the same partition or community are assigned shared gene identifiers.

### Read-to-isoform assignment

To quantify isoforms, labeled read paths must be assigned to compatible isoforms. Reads sharing identical splice structure (the same ordered sequence of splice-graph nodes) are aggregated into a single labeled read path with an associated read count. This aggregation is performed during Labeled Read Path Graph construction and enables efficient assignment by evaluating each unique splice pattern once rather than processing individual reads redundantly.

Isoforms (reconstructed or provided as reference annotations) are similarly assigned labeled paths through the splice graph based on their structural representation within the splice graph. Each labeled read path is compared against each isoform’s labeled read path to establish compatibility. Assignment proceeds through a cascading hierarchy of stringency levels, attempting the most stringent criteria first and progressively relaxing constraints until a compatible isoform is identified.

The assignment cascade operates in two phases reflecting HiFi and non-HiFi data characteristics. HiFi-style assignment (Phase 1) leverages high-accuracy boundary information by requiring TSS and PolyA nodes to match when present. If no assignment is made under HiFi rules, the algorithm falls back to non-HiFi-style assignment (Phase 2), trimming boundary nodes entirely to accommodate reads lacking reliable terminal information. Within each phase, multiple compatibility tests are applied in order of decreasing stringency.

### Phase 1: HiFi-style assignment (TSS/PolyA boundaries respected)

1. **Exact match with boundary anchoring:** Read path exactly matches isoform path end-to-end. If the read or isoform begins or ends with TSS or PolyA nodes, boundaries must match precisely.
2. **Compatible and contained with boundary anchoring:** Read path is fully contained within the isoform path as a consecutive segment. Boundaries must match when present.
3. **Introns contained with boundary anchoring:** All read introns appear in the isoform in the same order within the read’s genomic span, with no extra isoform introns intervening, and ≥75% read overlap with isoform structure. Boundaries must match when present.
4. **Compatible overlap with boundary anchoring:** Read and isoform are compatible (no conflicting introns) across their overlapping region, with no gaps in the overlap, and ≥75% read coverage. Boundaries must match when present.
5. **Compatible overlap without boundary anchoring:** Same as above, but TSS/PolyA boundaries need not match even if present.

### Phase 2: Non-HiFi-style assignment (TSS/PolyA boundaries ignored)

If no assignment is made in Phase 1, TSS and PolyA nodes are trimmed from both read and isoform paths, and compatibility tests are applied without boundary constraints:

6. **Exact match:** Trimmed paths match exactly.
7. **Compatible and contained:** Trimmed read path is fully contained in the isoform as a consecutive segment.
8. **Introns contained:** All read introns appear in the isoform in the same order within the r ad’s genomic span, with no extra isoform introns intervening, and ≥75% read overlap with isoform structure.
9. **Compatible overlap:** Read and isoform are compatible (no conflicting introns) across their overlapping region, with no gaps in the overlap, and ≥75% read coverage.

For labeled read paths compatible with multiple isoforms, assignment ambiguity is resolved probabilistically using relative expression levels estimated via EM (described below). Labeled read paths are additionally weighted by their 3′ end proximity to isoform 3′ termini to account for alternative polyadenylation, with weights decreasing linearly as genomic distance increases.

### Isoform quantification

Isoform abundances are estimated using an expectation-maximization (EM) algorithm that resolves ambiguous read assignments (building on EM-based frameworks described in (Bray et al., 2016; Kabza et al., 2024; B. Li & Dewey, 2011; Patro et al., 2017; Zare Jousheghani et al., 2025). Read-isoform compatibility follows the cascade described in ‘Read-to-Isoform Assignment’: the read’s labeled path must agree with the isoform’s exon-intron structure, with exact matches attempted first and progressively relaxed overlap tests (≥75% coverage) applied when needed. Reads are additionally weighted by their 3’ end proximity to isoform 3’ termini to account for alternative polyadenylation: For a read compatible with multiple isoforms, the weight assigned to each isoform index ***i*** is proportional to ***1 – di/D***, where *d_i_* is the genomic distance between the read’s 3’ end and isoform ***i***’s 3’ terminus, and ***D*** is the sum of distances across all compatible isoforms; explicitly, ***wri = 1 – di/D***. This linear weighting scheme prioritizes isoforms whose 3’ ends align more closely with the read’s observed termination point. When the ‘––oversimplify’ option is active for selected contigs, this EM stage is bypassed and reads are instead assigned to the single transcript with the greatest exonic base-pair overlap.

Let 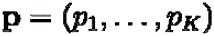 be the isoform proportions for gene 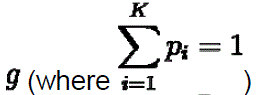, and let ***R_g_*** be the set of read compatibility classes (RCCs) assigned to gene ***g***. Each RCC 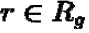 corresponds to a unique labeled read path and aggregates ***η_r_*** individual reads sharing that structure and assigned to the corresponding RCC. The likelihood of observing RCC ***r*** is:

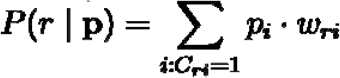

where 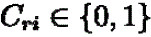 indicates compatibility between RCC ***r*** and isoform ***i***. To regularize low-support isoforms, a prior with base concentration **α = 0.01** is applied, with per-isoform prior mass scaled by the ambiguous RCC support assigned to that isoform (see below).

In the **expectation step**, posterior probabilities (i.e. fractions of each RCC assigned to each compatible isoform) are computed per RCC:

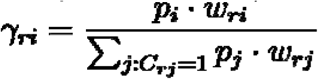

In the **maximization step**, isoform proportions (relative expression levels) are updated with scaling the prior by the amount of ambiguous RCC support assigned to each isoform. Let ***A_i_*** be the total read count summed across RCCs that map to more than one isoform but include isoform ***i***; then the effective prior mass is ***α_i_*** = ***α*** · ***A_i_***. The update becomes

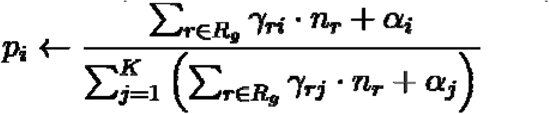

where ***η_r_*** is the total read count represented by RCC ***r*** and ***α_i_*** = **0** when isoform ***i*** has no ambiguous RCC support.

Iteration continues until the relative change in log-likelihood falls below **10^-6^** or maximum iterations are reached (250 for quantification-only mode, 1000 during assembly). From the final proportions, unique read counts (RCCs assigned exclusively to a single isoform with 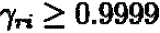), total fractional read counts, isoform fractions, and TPM are computed.

### LRAA single-cell RNA-seq automated workflow

LRAA-singlecell is implemented as a Workflow Description Language (WDL) workflow, enabling portable execution across cloud platforms, high-performance computing clusters, and local compute environments. Unless otherwise noted, all isoform discovery and quantification steps apply the LRAA algorithms described above, with parameters adapted for single-cell data characteristics as specified below.

### Input and preprocessing

LRAA-singlecell requires coordinate-sorted, indexed BAM files of long-read single-cell RNA-seq data aligned to a reference genome, the reference genome in FASTA format, and cell barcode annotations embedded in BAM read tags. Cell barcodes are read from a configurable BAM tag (default: CB, following the 10x Genomics convention), and unique molecular identifiers (UMIs) are read from a separate tag (default: XM).

For cluster-guided mode, the workflow additionally requires cell cluster assignments as input. These are provided as a tab-separated file mapping cell barcodes to cluster identifiers. Cluster assignments may be generated by the basic-mode pipeline or provided from external clustering analyses. Reference gene annotations in GTF format are optional for discovery modes but required for quantification-only operation.

### Basic mode workflow

#### Initial isoform discovery

In basic mode, LRAA first performs transcript isoform discovery on the complete dataset by pooling all single-cell reads into a unified pseudobulk analysis. This initial discovery step constructs a splice graph from all cell barcodes combined, treating the dataset as conventional bulk RNA-seq to establish an initial catalog of expressed isoforms across the entire cell population. During this pooled quantification, read tracking information is recorded for each aligned read, including the cell barcode, read name, assigned isoform(s), splice structure, and assignment confidence. This tracking file serves as the foundation for downstream single-cell count matrix construction.

#### Single-cell sparse matrix construction

For each unique cell barcode in the dataset, reads are aggregated at three hierarchical levels: (1) gene-level counts, summing all reads assigned to any isoform of a gene; (2) isoform-level counts, tallying reads assigned to specific transcript isoforms; and (3) splice pattern-level counts, grouping reads by unique splice junction combinations regardless of transcript identity. Splice patterns are encoded as hash codes derived from the ordered sequence of intron coordinates (as inspired by Isosceles (Kabza et al., 2024)), enabling detection of alternative splicing events independent of full-length isoform reconstruction.

#### Cell quality filtering

To distinguish genuine cells from empty droplets and low-quality cells, an optional filtering step applies the DropletUtils emptyDrops statistical framework (Lun et al., 2019). The gene-level sparse matrix is used to model the ambient RNA profile from low-count barcodes and to compute false discovery rates (FDR) for each barcode under the null hypothesis that observed counts arise solely from ambient contamination. Barcodes with FDR ≤ 0.01 (default) are retained as high-confidence cells.

#### Cell clustering

Cells are clustered using a Seurat (Hao et al., 2024) based workflow implemented in R that applies standard single-cell normalization, dimensionality reduction, and graph-based clustering to the filtered gene-level sparse matrix. The workflow follows conventional Seurat practices:

1. Genes expressed in fewer than a specified minimum number of cells (default: 10) are filtered to remove rare features.
2. Cells expressing fewer than a specified minimum number of genes (default: 1000) or exceeding a maximum threshold of mitochondrial content (default: 20%) are removed as low quality.
3. Raw counts are log-normalized to 10,000 reads per cell.
4. The top most highly variable genes (default: 2000) are identified via variance-stabilizing transformation.
5. Principal component analysis (PCA) is performed using the top variable genes, retaining the top principal components (default: 12).
6. A shared nearest neighbor (SNN) graph is constructed in PCA space and clustered using the Louvain algorithm (Blondel et al., 2008) at resolution (default: 0.6).
7. Uniform Manifold Approximation and Projection (UMAP) (McInnes et al., 2018) is computed for visualization.

Cell clustering results are exported as a tab-separated assignment file mapping each cell barcode to its cluster identifier. This file serves as input for the downstream cluster-guided LRAA workflow.

### LRAA cell cluster-guided mode workflow

#### BAM partitioning by cell cluster

Following cell clustering, either from basic mode or from provided precomputed assignments, the input BAM file is partitioned by cell cluster identity. Each read is examined for its cell barcode tag (default: CB) and matched against cluster assignments. Reads are written to cluster-specific BAM files (one per cluster). Unmapped reads, and reads from barcodes not present in the cluster assignments, are excluded. This partitioning produces a set of pseudobulk BAM files, each containing all reads from cells belonging to a specific cluster. BAM files are coordinate-sorted and indexed to enable efficient downstream processing. The partitioned BAM files are packaged into a compressed tarball for convenient retrieval and archival.

#### Per-cluster isoform discovery

Each cluster-specific BAM is processed independently through the complete LRAA isoform discovery pipeline. When the initial discovery GTF is provided (derived from basic mode or provided as a reference), it serves as an initial annotation for reference-guided LRAA isoform identification as applied to each cluster-specific BAM.

#### Cell cluster GTF merging for a consensus isoform catalog

Per-cluster GTF files are merged into a unified consensus annotation using a specialized execution of LRAA that treats the individual cluster-based isoform structures as collections of long RNA-seq reads, while retaining annotated TSS and PolyA sites. This merging procedure aggregates isoforms across cell clusters while resolving redundancy. Isoforms with identical exon–intron structures (identical splice junctions and overlapping terminal exons) are collapsed into single consensus transcripts, with gene assignments unified across clusters. Novel isoforms discovered in only one cluster are included in the final annotation if they pass expression and support thresholds, ensuring that well-supported cluster-specific variants are retained. The final merged GTF represents a comprehensive isoform catalog spanning all cell types and incorporates both shared isoforms expressed across clusters and cluster-specific transcript variants.

#### Final quantification and single-cell matrix generation

After generating the consensus GTF, each cluster of cells is re-quantified against the merged isoform catalog while leveraging an identical splice graph to produce final abundance estimates. This ensures that all cells are quantified using the same unified isoform catalog. Per-cluster read-to-isoform tracking files are merged into a single comprehensive tracking file spanning all cells and clusters, which serves as input for final sparse matrix construction. The workflow then yields separate sparse feature-by-cell count matrices at gene-, isoform-, and splice pattern-level resolution.

#### Reference annotation gene symbol integration

To facilitate interpretation and integration with reference annotations, an optional gene symbol incorporation step annotates LRAA-discovered isoforms with official gene symbols from a reference GTF. This procedure uses gffcompare (G. Pertea & Pertea, 2020) to compare the LRAA GTF to a reference annotation (e.g., GENCODE (Mudge et al., 2025)), identify matching or overlapping loci, and transfer gene symbols to LRAA transcripts based on structural similarity. Matching criteria include exact matches, contained matches, and intron-chain-compatible matches, with priority given to transcripts exhibiting high structural concordance. After gene symbol assignment, all single-cell sparse matrices (gene, isoform, and splice pattern levels) are updated to incorporate gene symbols into the LRAA-assigned feature identifiers (genes, isoforms, and splice patterns).

#### LRAA single cell workflow architecture and optimization

The workflow leverages parallel execution at multiple stages. Initial discovery can be parallelized across chromosomes or genomic contigs when targeted molecules are specified. LRAA isoform discovery is restricted to the nuclear genome. Mitochondrial genes are quantified based on reference gene annotations or simply quantified as reads per mitochondrial genome strand for estimating cellular mitochondrial read content in annotation-free mode. Per-cell-cluster discovery in cluster-guided mode executes clusters in parallel via WDL scatter blocks, and per-cluster quantification similarly parallelizes across cluster-specific BAMs. Resource allocation is configurable through workflow inputs, with separate memory and CPU parameters for discovery, quantification, clustering, and matrix-building tasks.

To support iterative analyses and avoid recomputation, the workflow accepts optional precomputed inputs. Users can provide a precomputed initial GTF and tracking files to skip initial discovery, or precomputed cluster assignments to skip cell filtering and clustering. This design enables expert users to provide curated cell cluster assignments for downstream cluster-guided isoform identification and quantification.

The workflow produces comprehensive outputs at each stage, including intermediate files for quality control and debugging. Outputs include initial and final GTF files, per-cluster BAMs and tracking files, pseudobulk expression matrices, sparse single-cell matrices, Seurat objects, UMAP visualizations, and gene symbol mappings. Cell type predictions were made using the Cellarium Cell Annotation Service (CAS) (Williams et al., 2025). A command-line utility is provided as part of the LRAA toolkit for automating submissions of cell gene expression sparse matrices to CAS for obtaining per-cell type predictions.

#### Single cell / nuclei analyses

Single cell and nuclei analyses were performed using Seurat v5 (Hao et al., 2024). Reference-only quantification was performed using LRAA in reference-guided mode along with GENCODE v47 reference annotations and minimap2 alignments of Kinnex reads to the GRCh38 reference genome. Gene-level sparse quantification matrices were loaded into Seurat and genes and cells were filtered requiring a minimum of 10 genes expressed per cell/nuclei, at least 500 genes expressed per cell/nuclei, and a maximum mitochondrial gene expression of 15% for cells and 5% for nuclei. Cell/nuclei biomarkers were identified using Seurat’s ‘FindAllMarkers’, and pairwise differential expression comparisons between cell/nuclei clusters using Seurat’s ‘FindMarkers’. Other Seurat analyses including generating UMAPs, clustering cells or nuclei and parameter settings are available in **Supplementary Code**. Functional enrichment analyses were performed using GProfiler (Kolberg et al., 2020).

#### Differential isoform usage testing

Differential isoform usage (DIU) was assessed by aggregating isoform-level read counts across all cells within each cluster to form pseudobulk matrices, followed by pairwise comparisons across every cluster pair; unspliced transcripts were excluded from all analyses. For splicing-focused analyses, isoforms sharing the same ordered intron chain were collapsed via splice-pattern hashcodes, summing their counts per cluster before DIU testing so that splice-junction usage changes are measured independently of transcript termini. For termini-focused analyses, isoforms were tested individually, and significant results were filtered to retain pairs that share splice patterns but differ in transcription start and/or end sites, isolating alternative start/stop usage.

Within each analysis mode, isoform (or splice-pattern group) usage fractions for a given gene were computed as

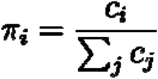

where ***c_i_*** is the pseudobulk read count for isoform (or group) ***i***. Usage differences between clusters A and B were summarized as

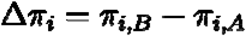

Isoforms were partitioned into dominant and alternate sets based on the magnitude and sign of **Δ*π_i_***, enabling detection of reciprocal isoform switches in which one set increases while the other decreases.

This framework adapts the pseudobulk strategy of Jogelkar et al. (2021) with several extensions: (1) reciprocal **Δ*π*** testing that requires both dominant and alternate sets to exceed effect-size thresholds (default **|Δ*π*|≥0.1**); (2) optional filtering on cell detection fraction so that, when a cell-fraction matrix is provided, each enriched isoform set must be detected in at least a specified fraction of cells (default 5%) in its enriched cluster; and (3) additional quality-control filters on minimum gene-level and isoform-set read depths (default ≥25 reads per cluster and per set). Genes passing these filters were evaluated by chi-squared contingency tests on 2×***N*** pseudobulk count tables, where rows correspond to clusters and columns to the tested isoforms/sets. P-values were adjusted with the Benjamini–Hochberg procedure, and DIU calls required both the FDR criterion (default FDR **<0.001**) and the effect-size thresholds described above.

#### Novel Proteoform Detection

We conducted a genome-wide search for proteoforms, focusing on transcripts classified as FSM, ISM, NIC, and NNIC. Open reading frames (ORFs) were predicted using TransDecoder v5.7.1 (*TransDecoder*, n.d.), and only complete ORFs containing both start and stop codons with minimum length thresholds of 50, 100, and 200 amino acids were retained for downstream analysis. Gene-specific reference splice junctions were extracted from the GENCODE v47 annotation (*GENCODE – Human Release 47*, n.d.). Novel splice junctions were identified by comparing transcript junction coordinates (donor-acceptor pairs) against the gene-specific reference junction set. Novel junctions where both the donor and acceptor sites fell within the CDS were classified as Novel Junctions. Intron Retention events were detected by identifying internal exons not present in the gene-specific reference annotation but utilizing known splice donor and acceptor sites for that gene. These internal exons represent retained intronic sequences. Retention events were considered to generate novel proteoforms if the retained intron overlapped with the CDS region.

To obtain protein-level counts, transcript counts encoding identical protein sequences were deduplicated. If any isoform encoding a given protein produced it without novel events (novel junctions or intron retention), the protein was classified as Reference Matched Proteoforms. Only proteins exclusively produced through novel splicing events were classified as Novel Proteoforms, with the event type (Novel Junction, Intron Retention, or both) determined by the randomly selected representative isoform.

The 3D structures of AIF1 proteoforms were modeled using ColabFold v1.5.5 (AlphaFold2 coupled with MMseqs2) with default parameters (Mirdita et al., 2022). Protein domains were predicted by InterProScan v5.76-107.0 (Jones et al., 2014). Structural superimposition and visualization were performed using UCSF ChimeraX v1.10.1 (Meng et al., 2023).

#### Generation of Simulated Long-Read Transcriptomic Data of Mouse and Arabidopsis

To establish a robust ground truth for benchmarking isoform identification and quantification tools, we generated simulated long-read datasets for mouse (*Mus musculus*) and Arabidopsis (*Arabidopsis thaliana*). Input transcript relative abundances were empirically derived from high-depth RNA-seq data (Mouse: SRR203276, (Grabherr et al., 2011); Arabidopsis: SRR3664432,(S. Li et al., 2016) and quantified using Salmon (v0.13.1) (Patro et al., 2017) against GENCODE vM32 (*GENCODE – Mouse Release M32*) and Araport11 (Cheng et al., 2017) annotations, respectively. These abundance profiles were normalized to a target sequencing depth of 20 million reads and integrated into a customized version of IsoSeqSim (v0.2) (available at (*IsoSeqSim at Devel_gpd_start*, n.d.), forked from (Y. Wang, n.d.)). This specific fork was modified to allow the simulator to initialize using a user-provided count matrix, ensuring simulated reads precisely reflected the empirical expression dynamics of the source datasets.

With IsoSeqSim, we simulated a gradient of four error profiles modeled on PacBio sequencing artifacts by adjusting the rates of substitutions (es), deletions (ed), and insertions (ei). This spanned a perfect control (0% error) and three error-prone datasets: low Error (1.6% total; es: 0.4%, ed: 0.6%, ei: 0.6%), medium Error (5.0% total; es: 1.7%, ed: 1.1%, ei: 2.2%), and high Error (8.5% total; es: 2.7%, ed: 1.8%, ei: 4.0%). Read incompleteness was modeled using IsoSeqSim-provided PacBio Sequel settings and dedicated parameters for 3’ (c3) and 5’ (c5) end completeness. Reads were mapped to their respective reference genomes (GRCm39 and TAIR10) using minimap2 (v2.28) (H. Li, 2018) with splice-aware presets (–ax splice) and processed with samtools (v1.15.1) (H. Li et al., 2009) into sorted BAM files.

To evaluate performance in reference-guided mode, we generated a reduced reference by randomly retaining exactly one isoform per gene from the full annotations. We then defined the expressed ground truth by filtering the original annotations for transcripts with non-zero abundance in our simulated data. By identifying the intersection between the single-isoform reduced reference and the expressed ground truth, we effectively partitioned the datasets into known isoforms (expressed and present in the reduced reference) and novel isoforms (expressed but artificially removed). This design allows for a rigorous evaluation of a tool’s ability to reconstruct missing isoforms while quantifying the whole transcriptome, by providing the reduced reference to the tools. Unlike the entire chromosome hold-out (Chen et al., 2023) or random isoform subsampling (Pardo-Palacios, Wang, et al., 2024), our approach creates a skeleton reference. By reducing transcript complexity while strictly maintaining gene loci, we were able to test how well discovery tools can reconstruct alternative splicing and novel isoforms against a constrained ground truth, rather than focusing on gene annotations.

#### Generation of Long-read Transcriptomic Data of SIRVs and Human Cell Lines

Four total RNA samples were leveraged to generate long-read transcriptomic data: Universal Human Reference RNA (UHRR), GM24385, K562, and BT474. cDNA synthesis and library preparation were performed following the standard PacBio Kinnex full-length RNA library protocol. For each cDNA synthesis reaction, 300 ng of RNA sample was spiked with 0.09 ng of one of the three SIRV Isoform Mixes (E0, E1, or E2) from SIRV-Set 1, targeting 1% of SIRV reads per RNA sample. cDNA synthesis reactions were performed in duplicate and cleaned with 3 serial 1.3X SMRTbell bead cleanups prior to amplification and concatenation. 24 final Kinnex libraries (comprising four distinct cell lines spiked with three different SIRV concentrations; each condition was performed in duplicate) were generated and pooled for sequencing. Each library was sequenced on the PacBio Revio system across 4 flow cells, yielding approximately 4 million reads per sample.

To evaluate isoform reconstruction accuracy using spike-in controls, deconcatenated HiFi poly(A) tail-trimmed reads were aligned to the GRCh38 reference genome supplemented with SIRV Set 1 sequences (https://www.lexogen.com/wp-content/uploads/2021/06/SIRV_Set1_Norm_Sequences_20210507.zip) using minimap2 (v2.28) (H. Li, 2018). Alignment was performed in splice-aware mode (–ax splice:hq). Post-alignment, reads were partitioned based on their mapping targets (SIRV contigs versus human ones), yielding sorted and indexed BAM files for independent evaluation. We defined the isoform quantification ground truth of spike-in by redistributing the total count of mapped SIRV reads across individual isoforms according to their manufacturer-defined stoichiometric ratios. Similar to the simulated datasets, we also generated benchmark references to evaluate reference-guided isoform discovery by parsing the GRCh38.gencode.v39 and SIRV annotation files to retain exactly one randomly selected transcript per gene.

### Generation of Long-read Transcriptomic Data of MORFs

#### MORF plasmid amplification and electroporation

Multiplexed Overexpression of Regulatory Factors (MORF) pooled plasmid library (comprising 1,836 genes encoded by 3,548 isoforms) was a gift from Feng Zhang (Addgene #1000000218) (Joung et al., 2023). Plasmid libraries were amplified by electroporation of 0.5 ug library into Endura electrocompetent cells (cat #60242-2) using BioRad Gene Pulser Xcell Total System (cat #1652660) with the following conditions: 2.5Kv: settings voltage 2500 V, capacitance 25 uF, resistance 200 Ω, cuvette 2 mm. Cells were immediately recovered by adding 1 mL of room temperature recovery media (cat #60242-2) and moving to a Celltreat recovery tube (cat #229475) and incubated on a shaker at 30°C for 20 minutes. Recovered cells were mixed into 500 ml LB media containing Carbenicillin (100ug/ml; from Broad Institute SQM) and incubated for 14 hours at 30°C. Serial dilutions were plated onto LB agar plates containing Carbenicillin (100ug/ml; from Broad Institute SQM) and incubated at 30°C for 18 hrs to quantify library diversity (55 million CFU). Plasmid purification was carried out using ZymoPURE II Plasmid Midiprep Kit (Zymo; D4201), splitting the 500 ml liquid cultures into eight separate purifications, followed by endotoxin removal (1800 ul at 276 ng/ul).

#### MORF plasmid QC via long read PacBio sequencing

To verify the integrity of the MORF plasmid library, full length MORFs were amplified off the plasmid DNA using primers flanking the EF1a promoter and WPRE containing “MARS” and “Venus” overhangs (DBO61 & DBO62; IDT) using the following conditions: 25 μl of 2x KAPA Hi-Fi Uracil ready mix (Roche), 5 ul of MORF plasmid library (50 ng/ul), 2.5 ul of 10 uM forward primerl (DBO61; IDT), 2.5 ul of 10 uM rev primer (DBO62; IDT), and 15 ul H20 to bring final volume to 50 ul. Reactions were amplified using the following cycling conditions: 98°C for 3 min, followed by 6 cycles (determined by qPCR) of 98°C for 20 s, 64°C for 30 s and 72°C for 8 min, followed by a final 72°C extension for 10 min. The amplified libraries were cleaned using 1.8× SPRIselect beads (Beckman Coulter B23318), eluted into 25 μl elution buffer and quantified with Qubit (Thermo Fisher Scientific, Q32851). The resulting amplicons were prepared for long-read sequencing using the PacBio 8-mer concatenation MAS-seq kit. Sequencing was performed on the PacBio Revio system.

#### MORF plasmid QC Analysis

PacBio HiFi reads were first segmented at adapter positions using skera (v1.2.0) and aligned to the MORF ORF insert reference (containing EF1a-ORF-Barcode-WPRE elements) using minimap2 (v2.28). To evaluate structural completeness, we examined minimap2 read alignments for corresponding reference transcript coverage. We further validated the sequences by performing a similarity search using BLAST+ (v2.15.0), querying the synthetic MORF sequences against the GENCODE v39 reference (*GENCODE – Human Release 39*, n.d.). We applied a stringent dual-threshold filter, retaining only those MORFs that exhibited both a sequence identity and a mean query coverage of >= 99%. This process retrieved 2,923 high-confidence isoforms from 1,566 human genes, providing a high-fidelity ground truth for the library prior to downstream experimental phases.

#### MORF transfection into HEK cells

HEK cells were transfected with 2.5 ug of MORF plasmid library using TransIT-LT1 Transfection Reagent (Mirus Bio; MIR 2304) following the manufacturer’s protocol. Briefly, 2.5 ug plasmid library was mixed into 250 ul OPTI-MEM and 7.5 ul TransIT-LT1 Reagent before incubating for 15 minutes at room temperature. Mixture was added to cells in a 6-well dish ∼80% confluency drop-wise to different areas in the well. Sample was gently rocked to evenly distribute the mix within complete media. To verify successful transfection, a plasmid expressing GFP (Addgene #145025; TFORF3549 GFP) was transfected in parallel in a separate well to check expression under a fluorescent microscope. Cells were harvested 72 hours post-transfection and total RNA was isolated using RNeasy Plus Mini Kit (Qiagen; cat # 74134) (1474 ng/ul in 60 ul water).

#### cDNA synthesis of MORF transcripts from HEK MORF total RNA

MORF transcripts (0.21 ul (300 ng) of HEK_MORF total RNA at 1474 ng/ul) were primed for reverse transcription using a custom primer annealing to the WPRE located on the 3’ end of the MORF transcripts (DBO140 [N14 UMI]; IDT) and cDNA synthesis was performed using Pacbio Iso-Seq express 2.0 kit (PacBio; cat no. 103-071-50) following manufacturer’s protocol. cDNA was prepared for PacBio sequencing using the Kinnex full-length RNA kit (PacBio; cat. no 103-072-000). Sequencing was performed on PacBio Revio system. The paired libraries before the concatenation were also sequenced on ONT MinION platform.

#### MORF data processing

To facilitate the analysis, a custom mini-genome of MORFs was constructed using the minigenome toolkit (*Minigenome: Makes a Mini-Genome, Shrinks Introns, Useful for Developing Small Test Sets and Small Data Sets for Workshops*, n.d.). The initial genome assembly was generated by shrinking introns and collapsing selected genes into single-contig representations based on the GRCh38 reference assembly and Gencode v39 annotations (*GENCODE – Human Release 39*, n.d.).

For ONT data, raw FASTQ reads were demultiplexed and trimmed after identifying 5’ and 3’ adapters based on Levenshtein distance (allowing for up to 1 mismatch) within a 150bp window. PacBio reads were demultiplexed at adapter sequences using skera (v1.2.0). Following demultiplexing, reads from both platforms were mapped to the MORF insert reference (EF1a-ORF-Barcode-WPRE) using minimap2 (v2.28) (H. Li, 2018). We disabled secondary alignments to ensure mapping uniqueness and applied a strict end-anchoring filtration step, retaining only primary, forward-strand alignments that extended exactly to the final nucleotide of the reference transcript to verify 3’-end completeness. Reads were assigned Transcript Names according to their corresponding reference sequence mappings.

Following initial alignment and filtering, soft-clipped bases were trimmed, and the final 14 nucleotides of each read were extracted as a UMI. To establish true molecular counts and remove PCR duplicates, reads were collapsed based on their assigned Transcript Name and UMI. This deduplication was performed by iterating through the alignments and retaining only the first primary alignment encountered for each unique UMI-Transcript Name pair; all subsequent reads sharing the same signature were discarded. Subsequently, this deduplicated dataset underwent a rigorous, systematic structural verification. We reconciled the deduplicated alignments with the MORF minigenome reference by extracting the intron chains and strand orientation for every retained read. A molecule was only included in the final benchmark if the read perfectly matched the annotated splice junction chain and strand orientation of its assigned isoform. Following the same strategy as our simulated mouse and Arabidopsis datasets, we derived two ground-truth reference sets from the MORF minigenome annotation: an Expressed Reference containing only isoforms with non-zero molecular counts, and a Reduced Reference with exactly one random isoform per gene.

### Generation of Long-read Transcriptomic Data for FTD Frontal Cortex Single Nuclei

#### Single nuclei isolation and cDNA library preparation from FTD postmortem frontal cortex

Postmortem medial frontal cortex brain tissue from a female donor with FTLD-TDP due to a progranulin mutation was obtained from the Mayo Clinic. On ice, approximately 45mg of tissue was homogenized using a Dounce homogenizer with 20 strokes of the loose pestle followed by 20 strokes of the tight pestle in 500uL chilled 1x Lysis Buffer 1 (10mM Tris-HCl, 10mM NaCl, 3mM MgCl2, 0.1% Nonidet P40 Substitute/IGEPAL CA-630, 1mM DTT and 1U/ul RNase inhibitors (Sigma-Aldrich, 03335402001)). Homogenate was incubated 4 minutes on ice then neutralized with 500uL chilled Wash Buffer (10mM Tris-HCl, 10mM NaCl, 3mM MgCl2, 1% Bovine Serum Albumin, 0.1% Tween-20, 1mM DTT). Large debris and cell clumps were removed by passing the suspension through a 70um Flowmi Cell Strainer. After lysis, nuclei were collected by centrifugation through a sucrose cushion gradient. Two sucrose gradient tubes were prepared by adding 500uL 1.8M Sucrose Cushion Buffer 1 (Sigma-Aldrich, NUC201-1KT), with the nuclei suspension carefully layered on top. Samples were then centrifuged at 16049 x g for 45 minutes at 4°C in a swinging bucket rotor. Myelin and debris were removed, and nuclei were resuspended in 500uL Wash Buffer. These were passed through a 40um Flowmi Cell Strainer, and aliquots were combined and centrifuged at 500 x g for 5 minutes at 4°C in a fixed angle rotor. The pellet was resuspended in 500uL PBS Buffer with 1% BSA and 1U/ul RNase inhibitors and counted using AO/PI on a Denovix CellDrop (Denovix, CD-AO-PI-1.5).

The resulting nuclei were processed using the 10x Genomics Chromium Next GEM Single Cell 3’ Reagents Kits v3.1 (10x Genomics, PN-1000268) and loaded on Chromium Next GEM Chip G (10x Genomics, 2000177) for targeted recovery of 10,000 nuclei. cDNA libraries were constructed as directed in the user guide (10x Genomics, CG000315 Rev. F). All libraries were quantified using Qubit dsDNA HS reagents (Invitrogen, Q33231). The average fragment size was determined using DNA HSD5000 screentape analysis (Agilent Technologies, 5067-5593, 5067-5592). cDNA libraries were between 400-2500 bp with an average size of 909bp. The resulting single nuclei cDNA libraries were then prepared for long read sequencing using Kinnex.

The library was sequenced across four PacBio flowcells and processed using the Kinnex Single-Cell workflow. Raw HiFi reads from each flowcell were de-concatenated to generate segmented reads (S-reads), with a consistently high proportion of full-length arrays across all flowcells (95.3–96.1%). Full-length non-concatemer (FLNC) reads were generated using the Iso-Seq refine and lima tools, during which poly(A) tails were removed, following PacBio’s recommended Iso-Seq processing guidelines.

#### Benchmarking Isoform Identification and Quantification

Benchmarking methods and related code were adapted from that used for the Isosceles paper (Kabza et al., 2024) (https://github.com/Genentech/Isosceles). As such, benchmarking is restricted to spliced isoforms where isoforms are uniquely identified according to their patterns of spliced introns. If multiple isoforms have shared splice patterns but differ at termini boundaries (alternative TSS or TES), they are collapsed into single representative isoforms and associated expression values or read counts are aggregated.

Isoform identification and quantification tools and software versions benchmarked included the following: LRAA v0.15.0, Bambu v3.4.0, ESPRESSO v1.5.0, FLAMES v0.1, FLAIR v2.0.0, Isosceles v0.2.0, IsoQuant v3.6.3, IsoSeq v4.0.0, Mandalorion v4.5.0, Oarfish v0.6.5, StringTie v2.2.3, and TALON v6.0.

**Isoform quantification accuracy**: Absolute relative differences (ARD) between estimated and truth set expression values (TPM) were computed like so:

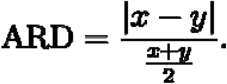

**Isoform identification accuracy**: Predicted isoforms are assigned as true positives (TP) based on matching reference isoform structures, false positives (FP) if missing from the reference annotation, or false negatives (FN) if present in the reference annotation but not predicted. Summary accuracy statistics true positive rate (TPR or aka. recall), positive predictive value (PPV aka. precision), and F1 scores are computed as follows:

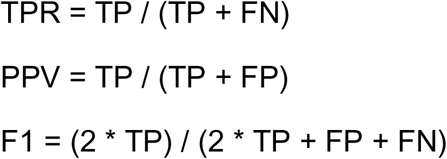

For annotation-free de novo isoform identification accuracy, reference annotations are not provided to prediction methods as input. All predicted isoforms are assigned TP, FP, or FN status based on comparison to the reference annotation truth set. For reference-guided isoform identification accuracy, predicted or reference isoforms corresponding to the splice patterns of those isoforms leveraged as guides were excluded from contributing to the benchmarking accuracy calculations.

Reference-guided isoform benchmarking experiments limited the reference isoform guide to a single isoform per gene.

When an exact truth set is not available, a proxy truth set is defined as the set of reference annotations having isoforms predicted by any one method in the corresponding evaluation. Isoforms predicted by multiple methods and not found within the reference annotation are ignored so that they do not contribute as TP or penalized as FP.

#### LRAA SQANTI-like read alignment and isoform structure annotation comparison

We incorporated a SQANTI (Pardo-Palacios, Arzalluz-Luque, et al., 2024) –like reference annotation comparison utility into the LRAA software suite to facilitate annotation of read alignments (bam) or isoform structures (GTF), considered ‘input features’, in comparison to a reference annotation. This SQANTI-like annotation utility was designed to be a flexible light-weight and efficient system for assigning input features to annotation categories based on references annotation splice pattern matching and genomic feature localization. Annotation categories are assigned separately for spliced and unspliced input features (**Supplementary Figure 5**).

#### Software availability

All source code, WDL workflows, and documentation are publicly available at https://github.com/MethodsDev/LongReadAlignmentAssembler under the MIT License.

## Acknowledgements

This work was supported by Broad Clinical Labs, a collaboration agreement between Pacific Biosciences of California, Inc. and the Broad Institute, and the Next Generation Fund Award from Broad Institute to AMA. MP was supported by the National Institutes of Health (R01NS120992, U54NS123743), the Target ALS Foundation, the BrightFocus Foundation (A2024017S), and the Kissick Family Foundation. CLT is the recipient of the Araminta Broch-Healey Endowed Chair in ALS. SA was the recipient of a Cullen Education and Research Foundation Young Investigator award and the Gluck ALS Research Scholar Award. CLT, MW and PB are supported by NIH/NINDS RM1NS133601 and the Chan Zuckerberg Initiative.

## Author contributions

BH, HY, and AMA designed the project. HY and BJH wrote the initial manuscript draft, performed analyses, and contributed to LRAA algorithm development. BJH led development of the LRAA software tool with contributions from HY, JW, CG, AK, and GA. DAB generated MORF data. AS and EW generated long-read RNA-seq libraries. XR, FH and SB prepared the single nuclei cDNA libraries. MP and DWD provided resources. VP advised on analysis and methods. AMA supervised the work. All authors contributed to the final manuscript.

## AI assistance disclosure

Generative AI tools (ChatGPT, Claude, Gemini) were used to assist with drafting and editing text and with assisting software development efforts. All AI-generated content was reviewed, tested, and revised by the authors, who take full responsibility for the final manuscript and software.

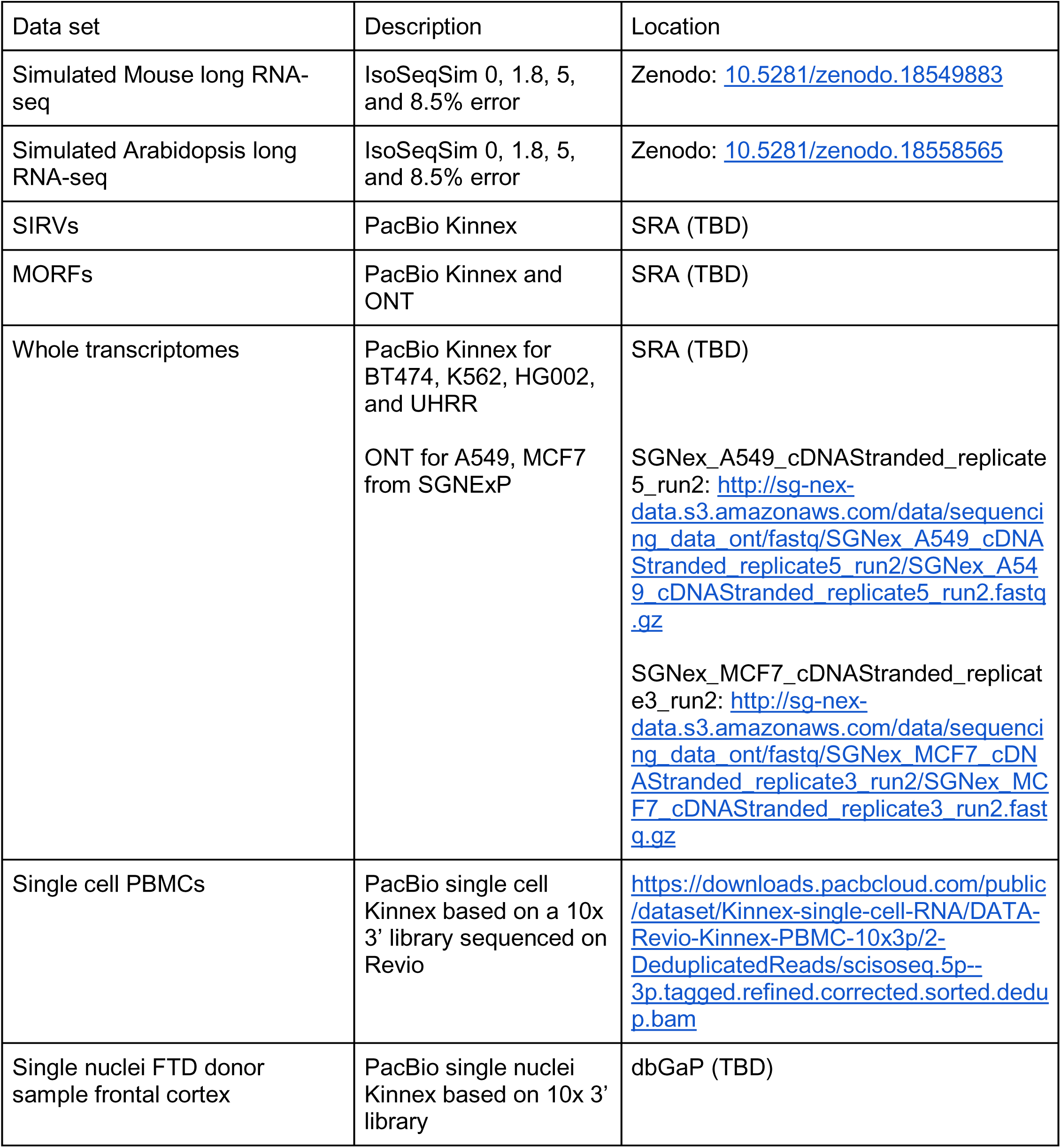
Data Resources Table.

## Supplementary Figures

**Supplementary Figure 1.**
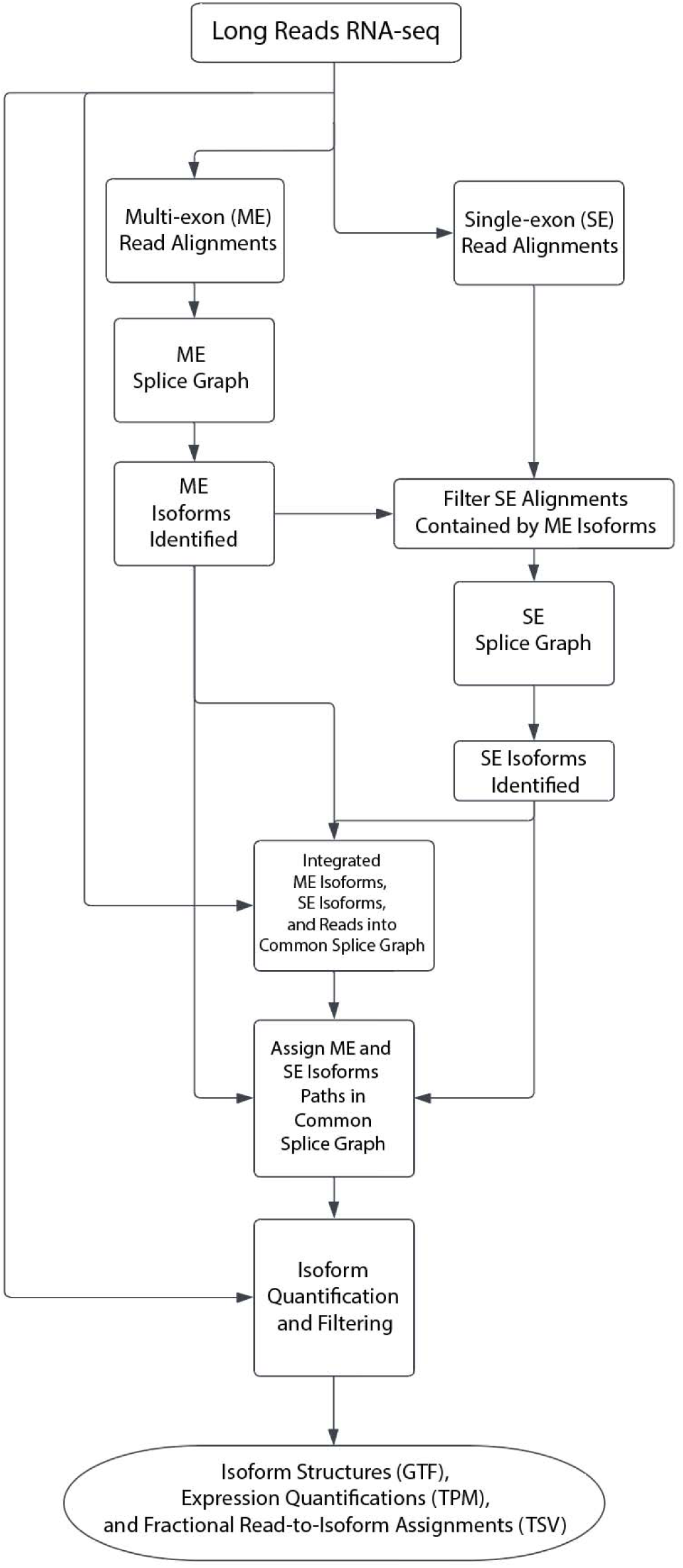
LRAA high-level algorithm for isoform identification and quantification starting from aligned long RNA-seq reads. See **Methods** for detailed descriptions of stages.

**Supplementary Figure 2:**
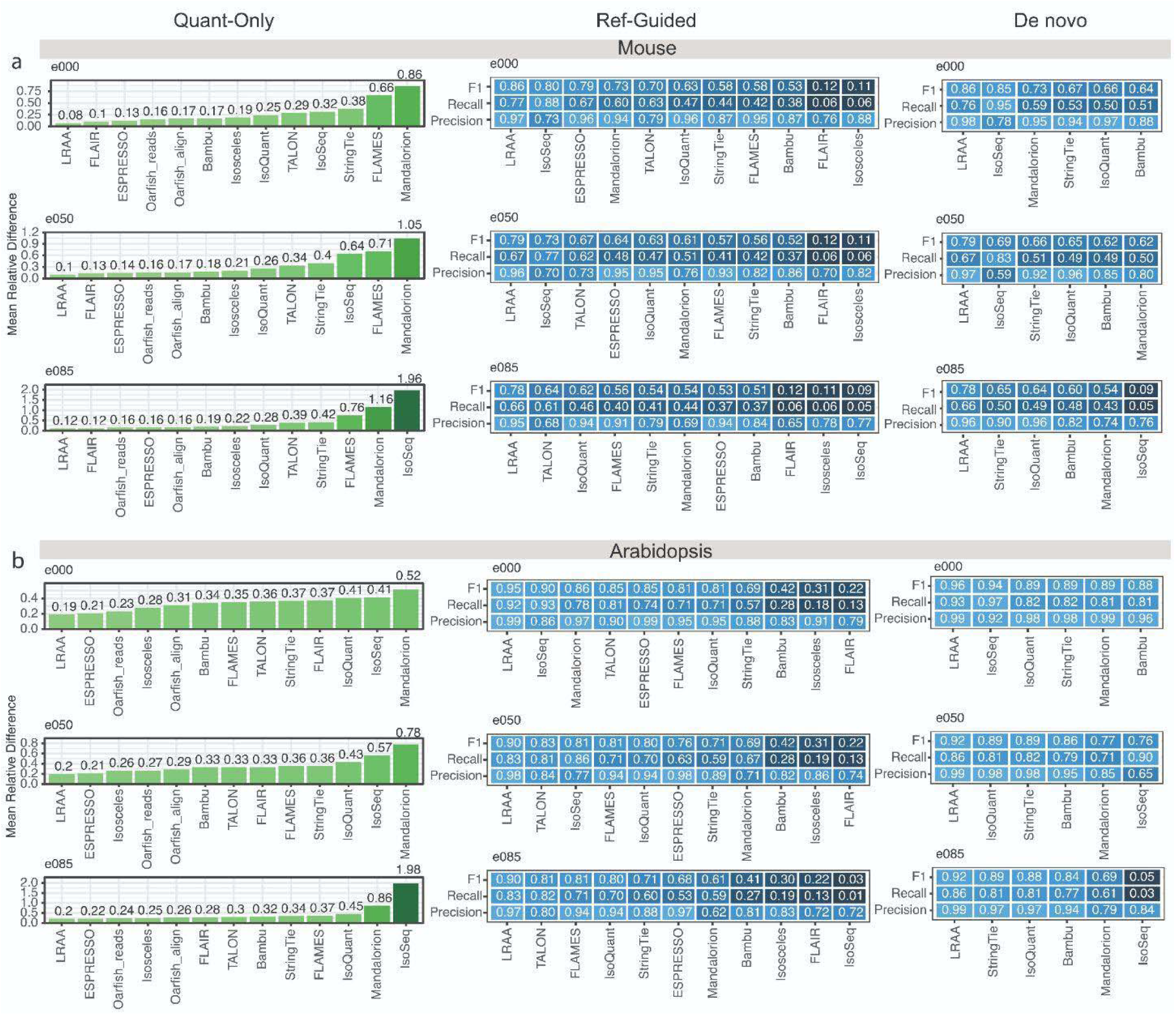
Isoform quantification and identification accuracy based on (a) Mouse or (b) Arabidopsis simulated long RNA-seq at varied levels of sequencing error: e000: error-free, e050: 5% error, and e085: 8.5% error.

**Supplementary Figure 3:**
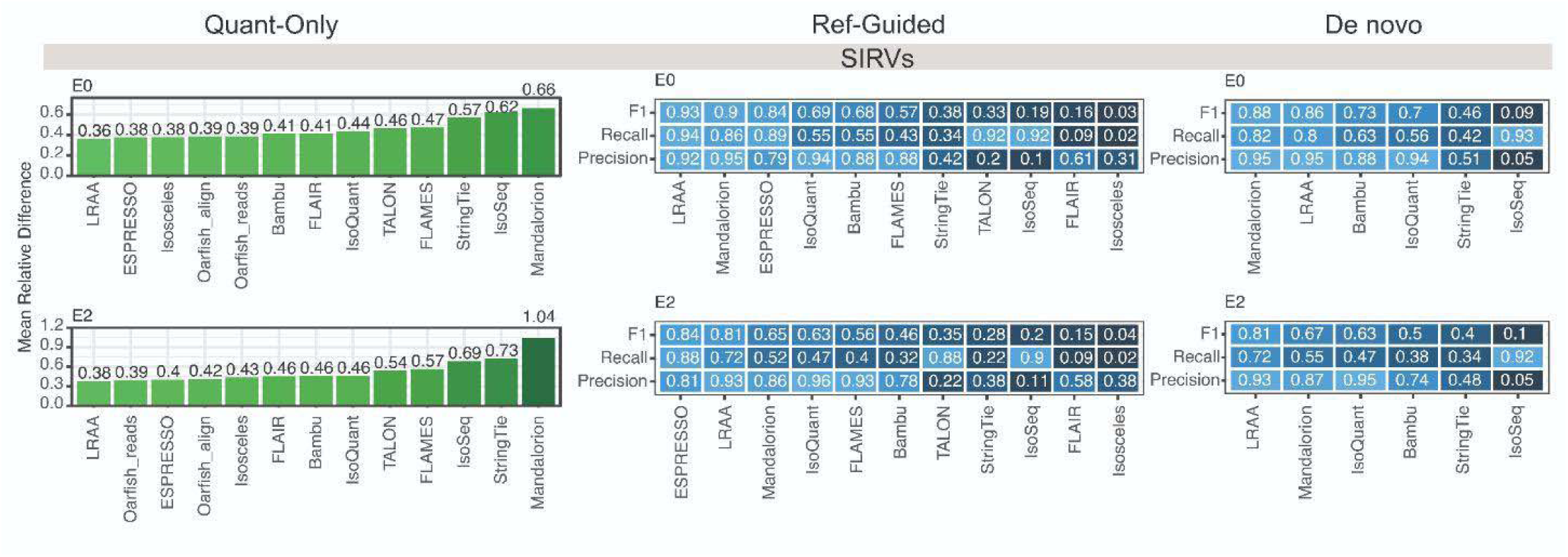
Isoform ID and Quant accuracy for SIRV mixes E0 and E2.

**Supplementary Figure 4:**
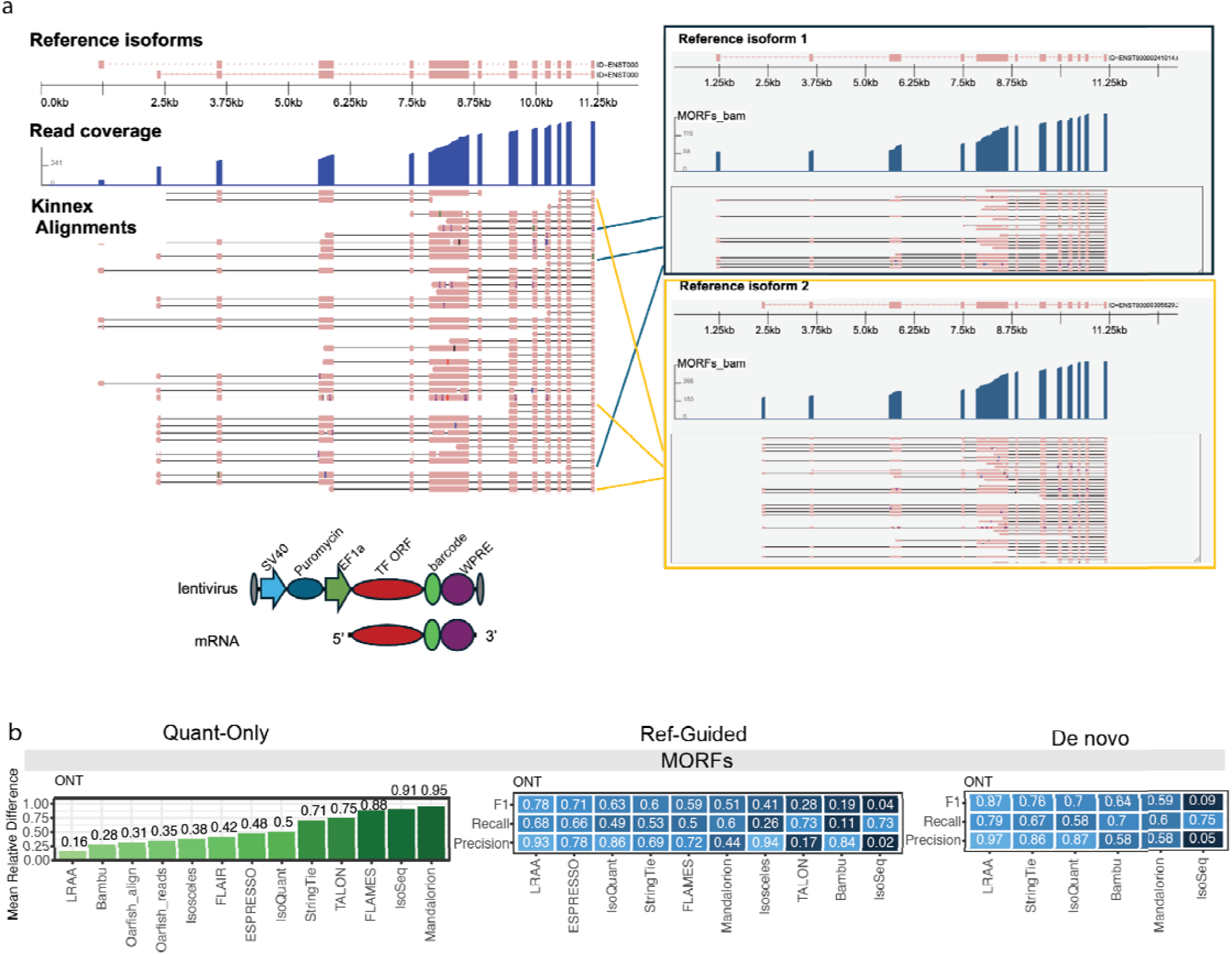
Isoform ID and Quantitation using MORFs. (a) Barcoding strategy enabling unambiguous identification of isoform from which read was derived, regardless of read-to-isoform structure mapping uncertainty. MORF lentivirus construct and mRNA structures shown at bottom, adapted from (Joung et al., 2023). (b) Quantification and isoform identification accuracy based on ONT long RNA-seq reads for MORFs.

**Supplementary Figure 5:**
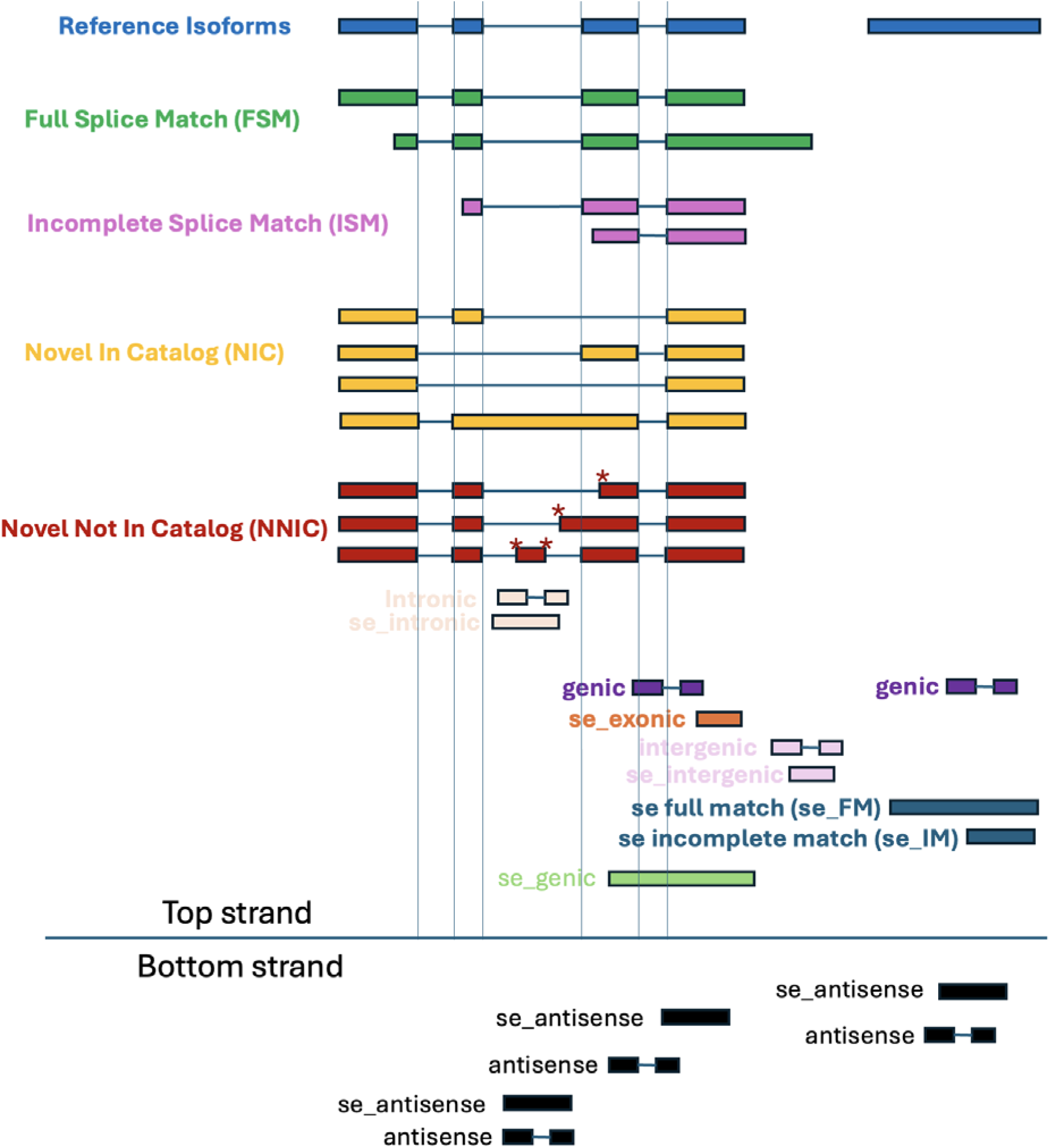
LRAA’s SQANTI-like classification scheme as applied to individual read alignments (BAM) or isoform structures (GTF) in comparison to a reference gene structure annotation (eg. GENCODE).

**Supplementary Figure 6:**
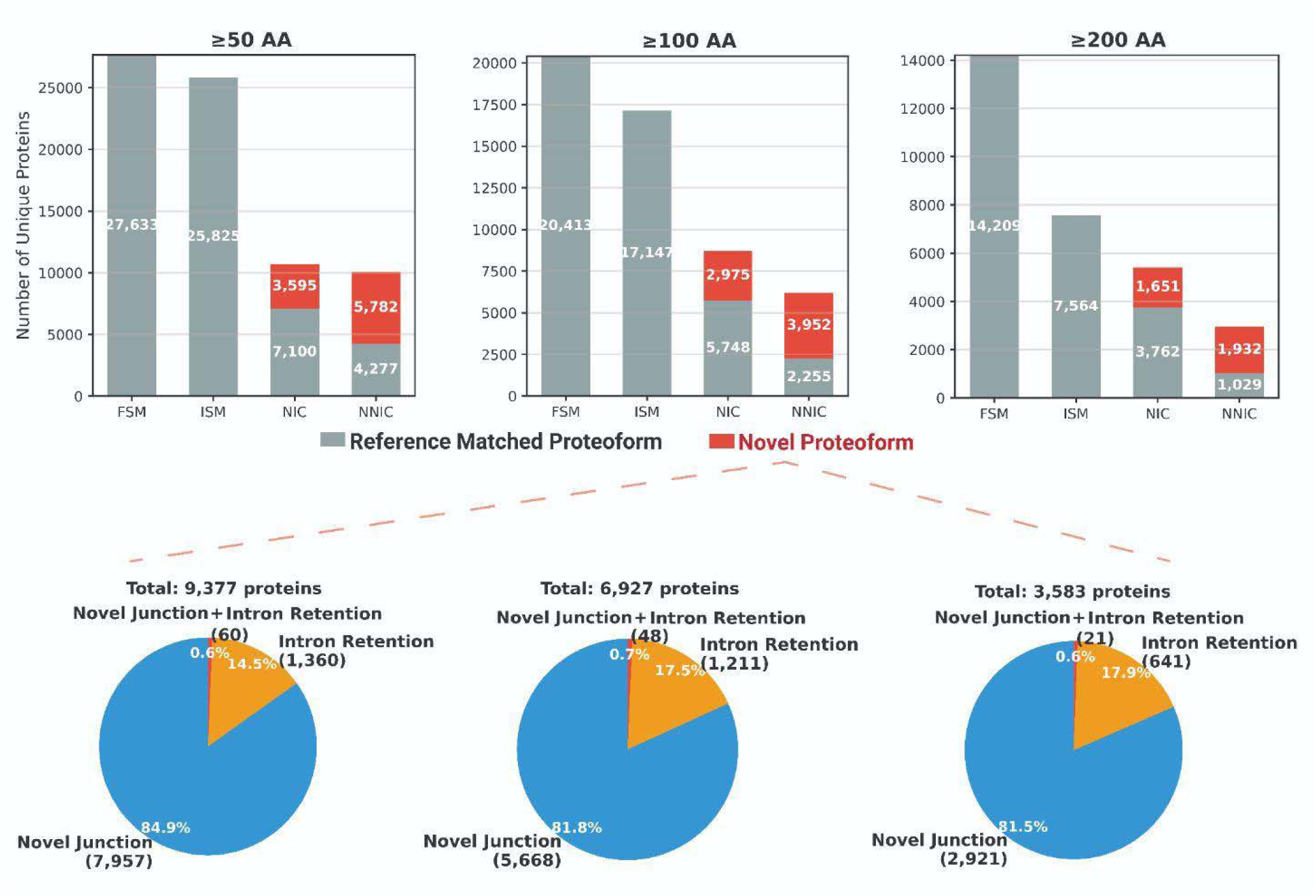
Novel proteoform identification in human PBMCs. Top: Stacked bar charts showing unique protein counts by classification (FSM, ISM, NIC, NNIC) at ≥50 AA, ≥100 AA, and ≥200 AA thresholds. Proteins classified as Reference-Matched Proteoforms (gray) or Novel Proteoforms (red) based on splice junction novelty or intron retention within coding sequences. Bottom: Pie charts showing novel proteoform composition (NIC + NNIC) by splicing event type: Novel Junction only (blue), Intron Retention only (gray), or both (orange).

**Supplementary Figure 7:**
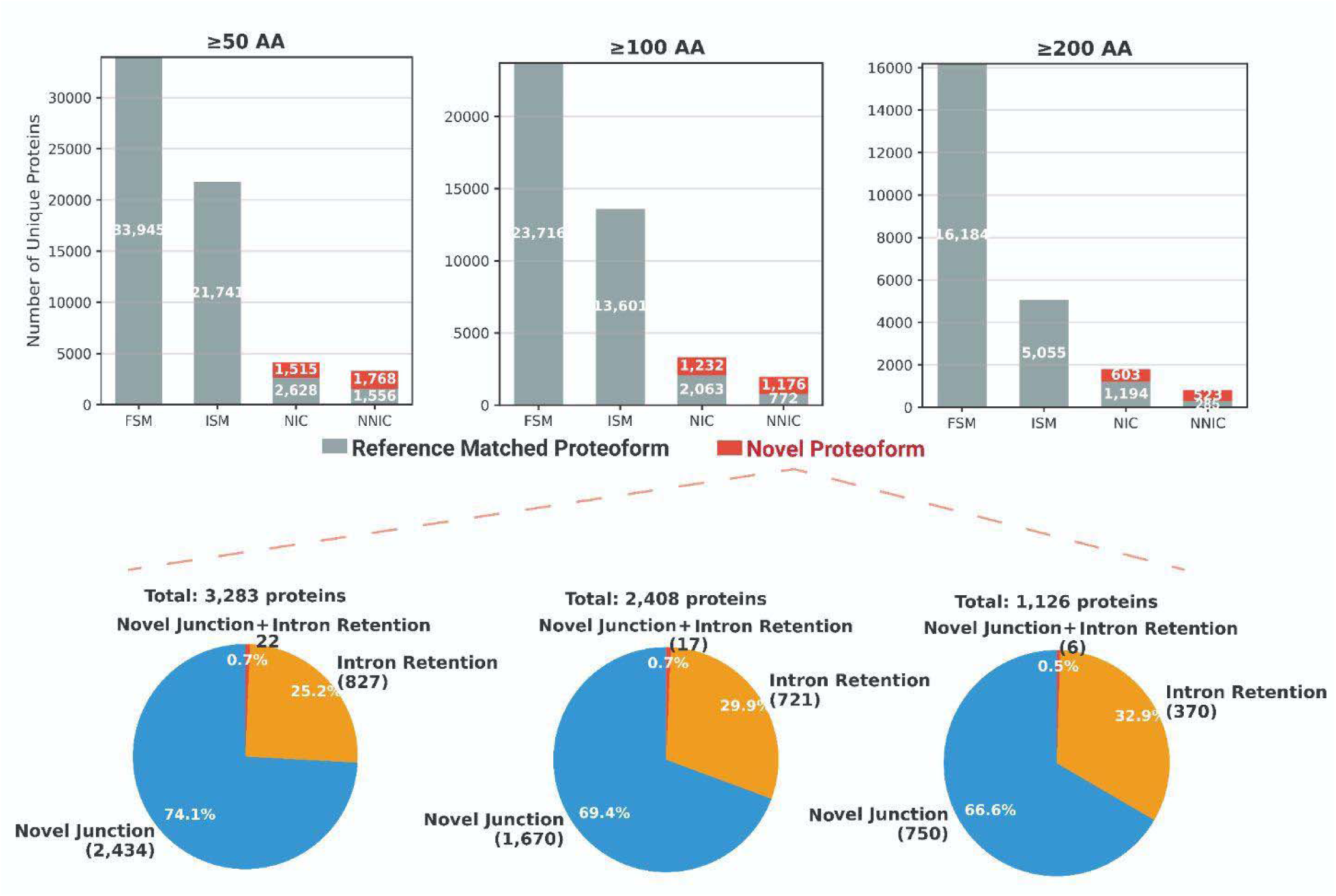
Novel proteoform identification in human FTD brain frontal cortex. Top: Stacked bar charts showing unique protein counts by classification (FSM, ISM, NIC, NNIC) at ≥50 AA, ≥100 AA, and ≥200 AA thresholds. Proteins classified as Reference-Matched Proteoforms (gray) or Novel Proteoforms (red) based on splice junction novelty or intron retention within coding sequences. Bottom: Pie charts showing novel proteoform composition (NIC + NNIC) by splicing event type: Novel Junction only (blue), Intron Retention only (gray), or both (orange).

## Notes

### Competing Interest Statement

The authors have declared no competing interest.

## References

1. Al’Khafaji, A. M., Smith, J. T., Garimella, K. V., Babadi, M., Popic, V., Sade-Feldman, M., Gatzen, M., Sarkizova, S., Schwartz, M. A., Blaum, E. M., Day, A., Costello, M., Bowers, T., Gabriel, S., Banks, E., Philippakis, A. A., Boland, G. M., Blainey, P. C., & Hacohen, N. (2024). High-throughput RNA isoform sequencing using programmed cDNA concatenation. Nature Biotechnology, 42(4), 582–586. 10.1038/s41587-023-01815-7

2. Amarasinghe, S. L., Su, S., Dong, X., Zappia, L., Ritchie, M. E., & Gouil, Q. (2020). Opportunities and challenges in long-read sequencing data analysis. Genome Biology, 21(1), 30. 10.1186/s13059-020-1935-5

3. Blondel, V. D., Guillaume, J.-L., Lambiotte, R., & Lefebvre, E. (2008). Fast unfolding of communities in large networks. In arXiv [physics.soc-ph]. arXiv. http://arxiv.org/abs/0803.0476

4. Bray, N. L., Pimentel, H., Melsted, P., & Pachter, L. (2016). Near-optimal probabilistic RNA-seq quantification. Nature Biotechnology, 34(5), 525–527. 10.1038/nbt.3519

5. Cheng, C.-Y., Krishnakumar, V., Chan, A. P., Thibaud-Nissen, F., Schobel, S., & Town, C. D. (2017). Araport11: a complete reannotation of the Arabidopsis thaliana reference genome. The Plant Journal: For Cell and Molecular Biology, 89(4), 789–804. 10.1111/tpj.13415

6. Chen, Y., Davidson, N. M., Wan, Y. K., Yao, F., Su, Y., Gamaarachchi, H., Sim, A., Patel, H., Low, H. M., Hendra, C., Wratten, L., Hakkaart, C., Sawyer, C., Iakovleva, V., Lee, P. L., Xin, L., Ng, H. E. V., Loo, J. M., Ong, X., … Göke, J. (2025). A systematic benchmark of Nanopore long-read RNA sequencing for transcript-level analysis in human cell lines. Nature Methods, 22(4), 801–812. 10.1038/s41592-025-02623-4

7. Chen, Y., Sim, A., Wan, Y. K., Yeo, K., Lee, J. J. X., Ling, M. H., Love, M. I., & Göke, J. (2023). Context-aware transcript quantification from long-read RNA-seq data with Bambu. Nature Methods, 20(8), 1187–1195. 10.1038/s41592-023-01908-w

8. Gao, Y., Wang, F., Wang, R., Kutschera, E., Xu, Y., Xie, S., Wang, Y., Kadash-Edmondson, K. E., Lin, L., & Xing, Y. (2023). ESPRESSO: Robust discovery and quantification of transcript isoforms from error-prone long-read RNA-seq data. Science Advances, 9(3), eabq5072. 10.1126/sciadv.abq5072

9. Garber, M., Grabherr, M. G., Guttman, M., & Trapnell, C. (2011). Computational methods for transcriptome annotation and quantification using RNA-seq. Nature Methods, 8(6), 469–477. 10.1038/nmeth.1613

10. *GENCODE – Human Release 39*. (n.d.). Retrieved February 6, 2026, from https://www.gencodegenes.org/human/release_39.html

11. *GENCODE – Human Release 47*. (n.d.). Retrieved February 6, 2026, from https://www.gencodegenes.org/human/release_47.html

12. *GENCODE – Mouse Release M32*. (n.d.). Retrieved February 6, 2026, from https://www.gencodegenes.org/mouse/release_M32.html

13. Glinos, D. A., Garborcauskas, G., Hoffman, P., Ehsan, N., Jiang, L., Gokden, A., Dai, X., Aguet, F., Brown, K. L., Garimella, K., Bowers, T., Costello, M., Ardlie, K., Jian, R., Tucker, N. R., Ellinor, P. T., Harrington, E. D., Tang, H., Snyder, M., … Cummings, B. B. (2022). Transcriptome variation in human tissues revealed by long-read sequencing. Nature, 608(7922), 353–359. 10.1038/s41586-022-05035-y

14. Grabherr, M. G., Haas, B. J., Yassour, M., Levin, J. Z., Thompson, D. A., Amit, I., Adiconis, X., Fan, L., Raychowdhury, R., Zeng, Q., Chen, Z., Mauceli, E., Hacohen, N., Gnirke, A., Rhind, N., di Palma, F., Birren, B. W., Nusbaum, C., Lindblad-Toh, K., … Regev, A. (2011). Full-length transcriptome assembly from RNA-Seq data without a reference genome. Nature Biotechnology, 29(7), 644–652. 10.1038/nbt.1883

15. Gupta, I., Collier, P. G., Haase, B., Mahfouz, A., Joglekar, A., Floyd, T., Koopmans, F., Barres, B., Smit, A. B., Sloan, S. A., Luo, W., Fedrigo, O., Ross, M. E., & Tilgner, H. U. (2018). Single-cell isoform RNA sequencing characterizes isoforms in thousands of cerebellar cells. Nature Biotechnology, 36(12), 1197–1202. 10.1038/nbt.4259

16. Gupta, P., O’Neill, H., Wolvetang, E. J., Chatterjee, A., & Gupta, I. (2024). Advances in single-cell long-read sequencing technologies. NAR Genomics and Bioinformatics, 6(2), lqae047. 10.1093/nargab/lqae047

17. Haas, B. J., Volfovsky, N., Town, C. D., Troukhan, M., Alexandrov, N., Feldmann, K. A., Flavell, R. B., White, O., & Salzberg, S. L. (2002). Full-length messenger RNA sequences greatly improve genome annotation. Genome Biology, 3(6), RESEARCH0029. 10.1186/gb-2002-3-6-research0029

18. Hao, Y., Stuart, T., Kowalski, M. H., Choudhary, S., Hoffman, P., Hartman, A., Srivastava, A., Molla, G., Madad, S., Fernandez-Granda, C., & Satija, R. (2024). Dictionary learning for integrative, multimodal and scalable single-cell analysis. Nature Biotechnology, 42(2), 293–304. 10.1038/s41587-023-01767-y

19. Hardwick, S. A., Hu, W., Joglekar, A., Fan, L., Collier, P. G., Foord, C., Balacco, J., Lanjewar, S., Sampson, M. M., Koopmans, F., Prjibelski, A. D., Mikheenko, A., Belchikov, N., Jarroux, J., Lucas, A. B., Palkovits, M., Luo, W., Milner, T. A., Ndhlovu, L. C., … Tilgner, H. U. (2022). Single-nuclei isoform RNA sequencing unlocks barcoded exon connectivity in frozen brain tissue. Nature Biotechnology, 40(7), 1082–1092. 10.1038/s41587-022-01231-3

20. Holmqvist, I., Bäckerholm, A., Tian, Y., Xie, G., Thorell, K., & Tang, K.-W. (2021). FLAME: long-read bioinformatics tool for comprehensive spliceome characterization. *RNA (New York*, N.Y.), 27(10), 1127–1139. 10.1261/rna.078800.121

21. *Iso-Seq Home*. (n.d.). Iso-Seq Docs. Retrieved February 10, 2026, from https://isoseq.how/

22. *IsoSeqSim at devel_gpd_start*. (n.d.). Github. Retrieved February 6, 2026, from https://github.com/MethodsDev/IsoSeqSim/tree/devel_gpd_start

23. Jones, P., Binns, D., Chang, H.-Y., Fraser, M., Li, W., McAnulla, C., McWilliam, H., Maslen, J., Mitchell, A., Nuka, G., Pesseat, S., Quinn, A. F., Sangrador-Vegas, A., Scheremetjew, M., Yong, S.-Y., Lopez, R., & Hunter, S. (2014). InterProScan 5: genome-scale protein function classification. *Bioinformatics (Oxford*, England), 30(9), 1236–1240. 10.1093/bioinformatics/btu031

24. Joung, J., Ma, S., Tay, T., Geiger-Schuller, K. R., Kirchgatterer, P. C., Verdine, V. K., Guo, B., Arias-Garcia, M. A., Allen, W. E., Singh, A., Kuksenko, O., Abudayyeh, O. O., Gootenberg, J. S., Fu, Z., Macrae, R. K., Buenrostro, J. D., Regev, A., & Zhang, F. (2023). A transcription factor atlas of directed differentiation. Cell, 186(1), 209–229.e26. 10.1016/j.cell.2022.11.026

25. Kabza, M., Ritter, A., Byrne, A., Sereti, K., Le, D., Stephenson, W., & Sterne-Weiler, T. (2024). Accurate long-read transcript discovery and quantification at single-cell, pseudo-bulk and bulk resolution with Isosceles. Nature Communications, 15(1), 7316. 10.1038/s41467-024-51584-3

26. Keren, H., Lev-Maor, G., & Ast, G. (2010). Alternative splicing and evolution: diversification, exon definition and function. Nature Reviews. Genetics, 11(5), 345–355. 10.1038/nrg2776

27. Klim, J. R., Williams, L. A., Limone, F., Guerra San Juan, I., Davis-Dusenbery, B. N., Mordes, D. A., Burberry, A., Steinbaugh, M. J., Gamage, K. K., Kirchner, R., Moccia, R., Cassel, S. H., Chen, K., Wainger, B. J., Woolf, C. J., & Eggan, K. (2019). ALS-implicated protein TDP-43 sustains levels of STMN2, a mediator of motor neuron growth and repair. Nature Neuroscience, 22(2), 167–179. 10.1038/s41593-018-0300-4

28. Kolberg, L., Raudvere, U., Kuzmin, I., Vilo, J., & Peterson, H. (2020). gprofiler2 –– an R package for gene list functional enrichment analysis and namespace conversion toolset g:Profiler. F1000Research, 9, 709. 10.12688/f1000research.24956.1

29. Kovaka, S., Zimin, A. V., Pertea, G. M., Razaghi, R., Salzberg, S. L., & Pertea, M. (2019). Transcriptome assembly from long-read RNA-seq alignments with StringTie2. Genome Biology, 20(1), 278. 10.1186/s13059-019-1910-1

30. Kumari, P., Kaur, M., Dindhoria, K., Ashford, B., Amarasinghe, S. L., & Thind, A. S. (2024). Advances in long-read single-cell transcriptomics. Human Genetics, 143(9-10), 1005–1020. 10.1007/s00439-024-02678-x

31. Li, B., & Dewey, C. N. (2011). RSEM: accurate transcript quantification from RNA-Seq data with or without a reference genome. BMC Bioinformatics, 12(1), 323. 10.1186/1471-2105-12-323

32. Li, H. (2018). Minimap2: pairwise alignment for nucleotide sequences. *Bioinformatics (Oxford*, England), 34(18), 3094–3100. 10.1093/bioinformatics/bty191

33. Li, H., Handsaker, B., Wysoker, A., Fennell, T., Ruan, J., Homer, N., Marth, G., Abecasis, G., Durbin, R., & 1000 Genome Project Data Processing Subgroup. (2009). The Sequence Alignment/Map format and SAMtools. Bioinformatics (Oxford, England), 25(16), 2078–2079. 10.1093/bioinformatics/btp352

34. Li, S., Yamada, M., Han, X., Ohler, U., & Benfey, P. N. (2016). High-resolution expression map of the Arabidopsis root reveals alternative splicing and lincRNA regulation. Developmental Cell, 39(4), 508–522. 10.1016/j.devcel.2016.10.012

35. Lun, A. T. L., Riesenfeld, S., Andrews, T., Dao, T. P., Gomes, T., participants in the 1st Human Cell Atlas Jamboree, & Marioni, J. C. (2019). EmptyDrops: distinguishing cells from empty droplets in droplet-based single-cell RNA sequencing data. Genome Biology, 20(1), 63. 10.1186/s13059-019-1662-y

36. Martin, J. A., & Wang, Z. (2011). Next-generation transcriptome assembly. Nature Reviews. Genetics, 12(10), 671–682. 10.1038/nrg3068

37. McInnes, L., Healy, J., Saul, N., & Großberger, L. (2018). UMAP: Uniform Manifold Approximation and Projection. Journal of Open Source Software, 3(29), 861. 10.21105/joss.00861

38. Melamed, Z. ‘ev, López-Erauskin, J., Baughn, M. W., Zhang, O., Drenner, K., Sun, Y., Freyermuth, F., McMahon, M. A., Beccari, M. S., Artates, J. W., Ohkubo, T., Rodriguez, M., Lin, N., Wu, D., Bennett, C. F., Rigo, F., Da Cruz, S., Ravits, J., Lagier-Tourenne, C., & Cleveland, D. W. (2019). Premature polyadenylation-mediated loss of stathmin-2 is a hallmark of TDP-43-dependent neurodegeneration. Nature Neuroscience, 22(2), 180–190. 10.1038/s41593-018-0293-z

39. Meng, E. C., Goddard, T. D., Pettersen, E. F., Couch, G. S., Pearson, Z. J., Morris, J. H., & Ferrin, T. E. (2023). UCSF ChimeraX: Tools for structure building and analysis. Protein Science, 32(11), e4792. 10.1002/pro.4792

40. Michielsen, L., Prjibelski, A. D., Foord, C., Hu, W., Jarroux, J., Hsu, J., Tomescu, A. I., Hajirasouliha, I., & Tilgner, H. U. (2025). Spatial isoform sequencing at sub-micrometer single-cell resolution reveals novel patterns of spatial isoform variability in brain cell types. In Neuroscience (No. biorxiv;2025.06.25.661563v2). bioRxiv. https://www.biorxiv.org/content/10.1101/2025.06.25.661563v1.full

41. *minigenome: makes a mini-genome, shrinks introns, useful for developing small test sets and small data sets for workshops*. (n.d.). Github. Retrieved February 6, 2026, from https://github.com/trinityrnaseq/minigenome

42. Mirdita, M., Schütze, K., Moriwaki, Y., Heo, L., Ovchinnikov, S., & Steinegger, M. (2022). ColabFold: making protein folding accessible to all. Nature Methods, 19(6), 679–682. 10.1038/s41592-022-01488-1

43. Mortazavi, A., Williams, B. A., McCue, K., Schaeffer, L., & Wold, B. (2008). Mapping and quantifying mammalian transcriptomes by RNA-Seq. Nature Methods, 5(7), 621–628. 10.1038/nmeth.1226

44. Mudge, J. M., Carbonell-Sala, S., Diekhans, M., Martinez, J. G., Hunt, T., Jungreis, I., Loveland, J. E., Arnan, C., Barnes, I., Bennett, R., Berry, A., Bignell, A., Cerdán-Vélez, D., Cochran, K., Cortés, L. T., Davidson, C., Donaldson, S., Dursun, C., Fatima, R., … Frankish, A. (2025). GENCODE 2025: reference gene annotation for human and mouse. Nucleic Acids Research, 53(D1), D966–D975. 10.1093/nar/gkae1078

45. Nilsen, T. W., & Graveley, B. R. (2010). Expansion of the eukaryotic proteome by alternative splicing. Nature, 463(7280), 457–463. 10.1038/nature08909

46. Pardo-Palacios, F. J., Arzalluz-Luque, A., Kondratova, L., Salguero, P., Mestre-Tomás, J., Amorín, R., Estevan-Morió, E., Liu, T., Nanni, A., McIntyre, L., Tseng, E., & Conesa, A. (2024). SQANTI3: curation of long-read transcriptomes for accurate identification of known and novel isoforms. Nature Methods, 21(5), 793–797. 10.1038/s41592-024-02229-2

47. Pardo-Palacios, F. J., Wang, D., Reese, F., Diekhans, M., Carbonell-Sala, S., Williams, B., Loveland, J. E., De María, M., Adams, M. S., Balderrama-Gutierrez, G., Behera, A. K., Gonzalez Martinez, J. M., Hunt, T., Lagarde, J., Liang, C. E., Li, H., Meade, M. J., Moraga Amador, D. A., Prjibelski, A. D., … Brooks, A. N. (2024). Systematic assessment of long-read RNA-seq methods for transcript identification and quantification. Nature Methods, 21(7), 1349–1363. 10.1038/s41592-024-02298-3

48. Patro, R., Duggal, G., Love, M. I., Irizarry, R. A., & Kingsford, C. (2017). Salmon provides fast and bias-aware quantification of transcript expression. Nature Methods, 14(4), 417–419. 10.1038/nmeth.4197

49. Pertea, G., & Pertea, M. (2020). GFF utilities: GffRead and GffCompare. F1000Research, 9, 304. 10.12688/f1000research.23297.2

50. Pertea, M., Shumate, A., Pertea, G., Varabyou, A., Breitwieser, F. P., Chang, Y.-C., Madugundu, A. K., Pandey, A., & Salzberg, S. L. (2018). CHESS: a new human gene catalog curated from thousands of large-scale RNA sequencing experiments reveals extensive transcriptional noise. Genome Biology, 19(1), 208. 10.1186/s13059-018-1590-2

51. Prjibelski, A. D., Mikheenko, A., Joglekar, A., Smetanin, A., Jarroux, J., Lapidus, A. L., & Tilgner, H. U. (2023). Accurate isoform discovery with IsoQuant using long reads. Nature Biotechnology, 41(7), 915–918. 10.1038/s41587-022-01565-y

52. Prudencio, M., Humphrey, J., Pickles, S., Brown, A.-L., Hill, S. E., Kachergus, J. M., Shi, J., Heckman, M. G., Spiegel, M. R., Cook, C., Song, Y., Yue, M., Daughrity, L. M., Carlomagno, Y., Jansen-West, K., de Castro, C. F., DeTure, M., Koga, S., Wang, Y.-C., … Petrucelli, L. (2020). Truncated stathmin-2 is a marker of TDP-43 pathology in frontotemporal dementia. The Journal of Clinical Investigation, 130(11), 6080–6092. 10.1172/JCI139741

53. Reese, F., Williams, B., Balderrama-Gutierrez, G., Wyman, D., Çelik, M. H., Rebboah, E., Rezaie, N., Trout, D., Razavi-Mohseni, M., Jiang, Y., Borsari, B., Morabito, S., Liang, H. Y., McGill, C. J., Rahmanian, S., Sakr, J., Jiang, S., Zeng, W., Carvalho, K., … Mortazavi, A. (2023). The ENCODE4 long-read RNA-seq collection reveals distinct classes of transcript structure diversity. In bioRxivorg. 10.1101/2023.05.15.540865

54. Sharon, D., Tilgner, H., Grubert, F., & Snyder, M. (2013). A single-molecule long-read survey of the human transcriptome. Nature Biotechnology, 31(11), 1009–1014. 10.1038/nbt.2705

55. Sim, A. D., Ling, M. H., Chen, Y., Lu, H., See, Y. X., Perrin, A., Leng Agnes, O. B., Cao, E. Y., Chia, B., Liu, J., Wüstefeld, T., Shin, J., & Göke, J. (2025). Isoform-level discovery, quantification and fusion analysis from single-cell and spatial long-read RNA-seq data with Bambu-Clump. In bioRxiv. 10.1101/2024.12.30.630828

56. Stark, R., Grzelak, M., & Hadfield, J. (2019). RNA sequencing: the teenage years. Nature Reviews. Genetics, 20(11), 631–656. 10.1038/s41576-019-0150-2

57. Steijger, T., Abril, J. F., Engström, P. G., Kokocinski, F., RGASP Consortium, Hubbard, T. J., Guigó, R., Harrow, J., & Bertone, P. (2013). Assessment of transcript reconstruction methods for RNA-seq. Nature Methods, 10(12), 1177–1184. 10.1038/nmeth.2714

58. Tang, A. D., Soulette, C. M., van Baren, M. J., Hart, K., Hrabeta-Robinson, E., Wu, C. J., & Brooks, A. N. (2020). Full-length transcript characterization of SF3B1 mutation in chronic lymphocytic leukemia reveals downregulation of retained introns. Nature Communications, 11(1), 1438. 10.1038/s41467-020-15171-6

59. Tardaguila, M., de la Fuente, L., Marti, C., Pereira, C., Pardo-Palacios, F. J., Del Risco, H., Ferrell, M., Mellado, M., Macchietto, M., Verheggen, K., Edelmann, M., Ezkurdia, I., Vazquez, J., Tress, M., Mortazavi, A., Martens, L., Rodriguez-Navarro, S., Moreno-Manzano, V., & Conesa, A. (2018). SQANTI: extensive characterization of long-read transcript sequences for quality control in full-length transcriptome identification and quantification. Genome Research, 28(3), 396–411. 10.1101/gr.222976.117

60. Teng, M., Love, M. I., Davis, C. A., Djebali, S., Dobin, A., Graveley, B. R., Li, S., Mason, C. E., Olson, S., Pervouchine, D., Sloan, C. A., Wei, X., Zhan, L., & Irizarry, R. A. (2016). A benchmark for RNA-seq quantification pipelines. Genome Biology, 17(1), 74. 10.1186/s13059-016-0940-1

61. Tilgner, H., Grubert, F., Sharon, D., & Snyder, M. P. (2014). Defining a personal, allele-specific, and single-molecule long-read transcriptome. Proceedings of the National Academy of Sciences of the United States of America, 111(27), 9869–9874. 10.1073/pnas.1400447111

62. *TransDecoder*. (n.d.). Github. https://github.com/TransDecoder/TransDecoder/wiki/Home

63. Volden, R., Palmer, T., Byrne, A., Cole, C., Schmitz, R. J., Green, R. E., & Vollmers, C. (2018). Improving nanopore read accuracy with the R2C2 method enables the sequencing of highly multiplexed full-length single-cell cDNA. Proceedings of the National Academy of Sciences of the United States of America, 115(39), 9726–9731. 10.1073/pnas.1806447115

64. Volden, R., Schimke, K. D., Byrne, A., Dubocanin, D., Adams, M., & Vollmers, C. (2023). Identifying and quantifying isoforms from accurate full-length transcriptome sequencing reads with Mandalorion. Genome Biology, 24(1), 167. 10.1186/s13059-023-02999-6

65. Wang, C., Prawer, Y. D. J., Voogd, O., Schuster, J., De Paoli-Iseppi, R., Li, A., Hallab, J., Tian, L., Peng, H., David, M., Du, M. R. M., Velasco, S., Garone, M. G., Dong, X., Zeglinski, K., Pavan, C., Law, K. C. L., Abu-Bonsrah, K. D., Hunt, C. P. J., … You, Y. (2025). Igniting full-length isoform analysis in single-cell and spatial RNA-seq data with FLAMESv2. In bioRxiv. 10.1101/2025.10.19.683327

66. Wang, Y. (n.d.). *IsoSeqSim: Iso-Seq reads simulator for PacBio and ONT full-length isoform sequencing technologies*. Github. Retrieved February 6, 2026, from https://github.com/yunhaowang/IsoSeqSim

67. Wang, Z., Gerstein, M., & Snyder, M. (2009). RNA-Seq: a revolutionary tool for transcriptomics. Nature Reviews. Genetics, 10(1), 57–63. 10.1038/nrg2484

68. Williams, S. R., Grab, F., Kamath, G. M., Ordabayev, Y., Mellen, J., Roelli, P., Cibulskis, K., Lehnert, E., Xie, F., Covarrubias, M., Rahman, N.-T., Tickle, T., Erhan, E., Malfroy-Camine, N., Lydon, K., Babadi, M., & Delaney, N. F. (2025). Accelerating scRNA-seq Analysis: Automated cell type annotation using representation learning and vector search. In Bioinformatics (No. biorxiv;2025.10.06.680787v1). bioRxiv. https://www.biorxiv.org/content/10.1101/2025.10.06.680787v1

69. Workman, R. E., Myrka, A. M., Wong, G. W., Tseng, E., Welch, K. C., Jr, & Timp, W. (2018). Single-molecule, full-length transcript sequencing provides insight into the extreme metabolism of the ruby-throated hummingbird Archilochus colubris. GigaScience, 7(3), 1–12. 10.1093/gigascience/giy009

70. Workman, R. E., Tang, A. D., Tang, P. S., Jain, M., Tyson, J. R., Razaghi, R., Zuzarte, P. C., Gilpatrick, T., Payne, A., Quick, J., Sadowski, N., Holmes, N., de Jesus, J. G., Jones, K. L., Soulette, C. M., Snutch, T. P., Loman, N., Paten, B., Loose, M., … Timp, W. (2019). Nanopore native RNA sequencing of a human poly(A) transcriptome. Nature Methods, 16(12), 1297–1305. 10.1038/s41592-019-0617-2

71. Wyman, D., Balderrama-Gutierrez, G., Reese, F., Jiang, S., Rahmanian, S., Forner, S., Matheos, D., Zeng, W., Williams, B., Trout, D., England, W., Chu, S.-H., Spitale, R. C., Tenner, A. J., Wold, B. J., & Mortazavi, A. (2019). A technology-agnostic long-read analysis pipeline for transcriptome discovery and quantification. In Genomics (No. biorxiv;672931v2). bioRxiv. https://www.biorxiv.org/content/10.1101/672931v2

72. Xu, J., Su, Z., Hong, H., Thierry-Mieg, J., Thierry-Mieg, D., Kreil, D. P., Mason, C. E., Tong, W., & Shi, L. (2014). Cross-platform ultradeep transcriptomic profiling of human reference RNA samples by RNA-Seq. Scientific Data, 1(1), 140020. 10.1038/sdata.2014.20

73. Zare Jousheghani, Z., Singh, N. P., & Patro, R. (2025). Oarfish: enhanced probabilistic modeling leads to improved accuracy in long read transcriptome quantification. *Bioinformatics (Oxford*, England), 41(Supplement_1), i304–i313. 10.1093/bioinformatics/btaf240

